## Supplemental Text for "The impacts of fine-tuning, phylogenetic distance, and sample size on big-data bioacoustics"

Short Title: Machine learning in bioacoustics

Kaiya L. Provost[1*], Jiaying Yang[1], Bryan C. Carstens[1]
1: Department of Evolution, Ecology and Organismal Biology, The Ohio State University. 318 W. 12th Ave. 300 Aronoff Laboratory Columbus, OH 43210

*: Corresponding author: Kaiya L. Provost,, Department of Evolution, Ecology and Organismal Biology, The Ohio State University. 318 W. 12th Ave. 300 Aronoff Laboratory Columbus, OH 43210

### Supplementary Figures

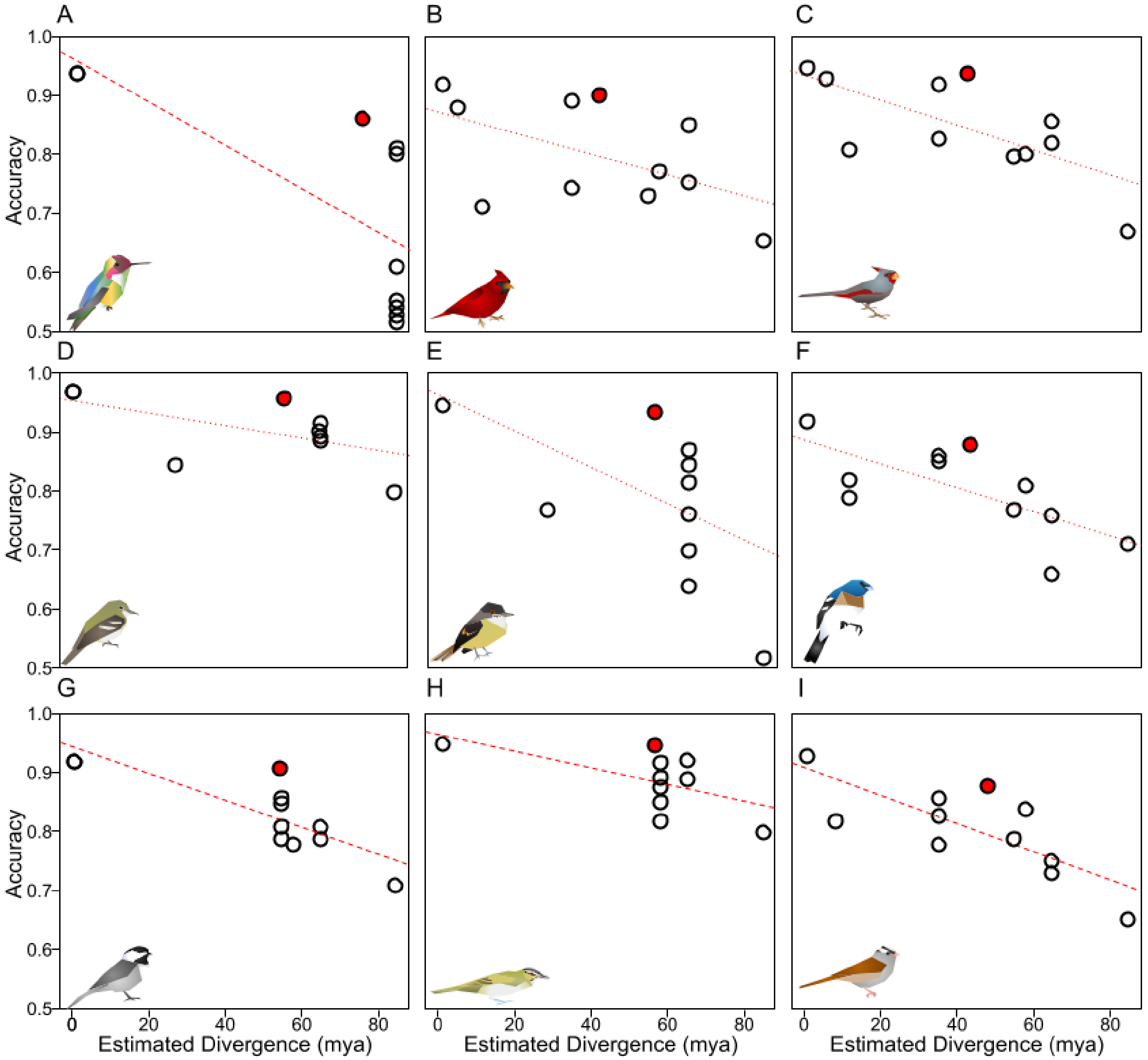

**S1 Fig: Model performance by species drops with estimated divergence, but training on multiple species performs better than expected given average divergence time**. X-axis gives the estimated divergence time. Y-axis gives the accuracy. Colored lines show the line of best fit, with solid lines bing significant and dotted lines not significant. Filled red points were trained on the “9 Species” model. A) *Calypte anna*. B) *Cardinalis cardinalis*. C) *Cardinalis sinuatus*. D) *Empidonax virescens*. E) *Myiarchus tuberculifer*. F) *Passerina amoena*. G) *Poecile carolinensis*. H) *Vireo altiloquus*. I) *Zonotrichia leucophrys*.

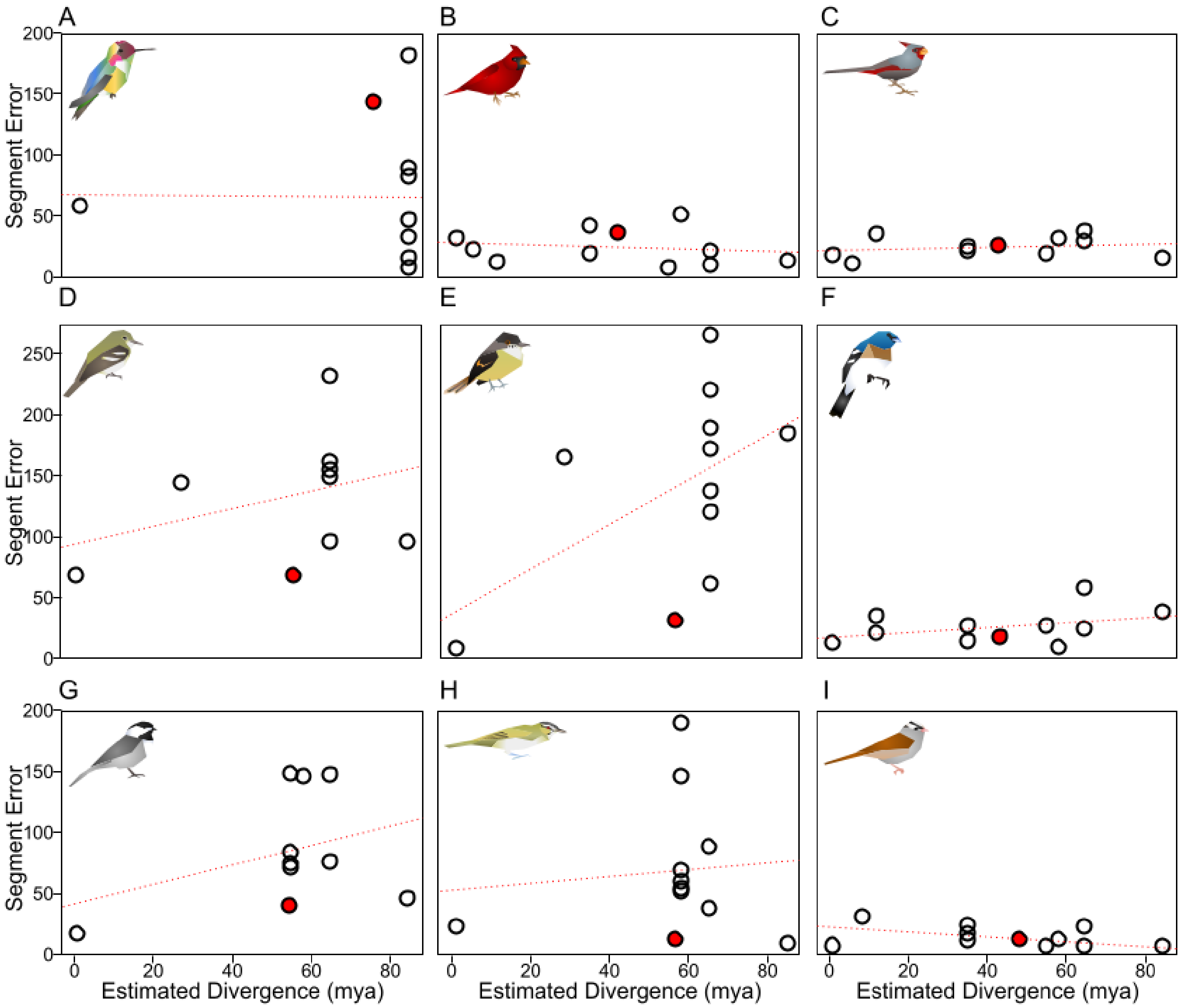

**S2 Fig: Segment error rate is worse with more distant divergence times, but training on multiple species performs better than expected**. X-axis gives the estimated divergence time. Y-axis gives the accuracy. Colored lines show the line of best fit, with solid lines being significant and dotted lines not significant. Filled red points were trained on the "9 Species" model. A) *Calypte anna*. B) *Cardinalis cardinalis.* C) *Cardinalis sinuatus.* D) *Empidonax virescens*. E) *Myiarchus tuberculifer*. F) *Passerina amoena.* G) *Poecile carolinensis.* H) *Vireo altiloquus.* I) *Zonotrichia leucophrys.*

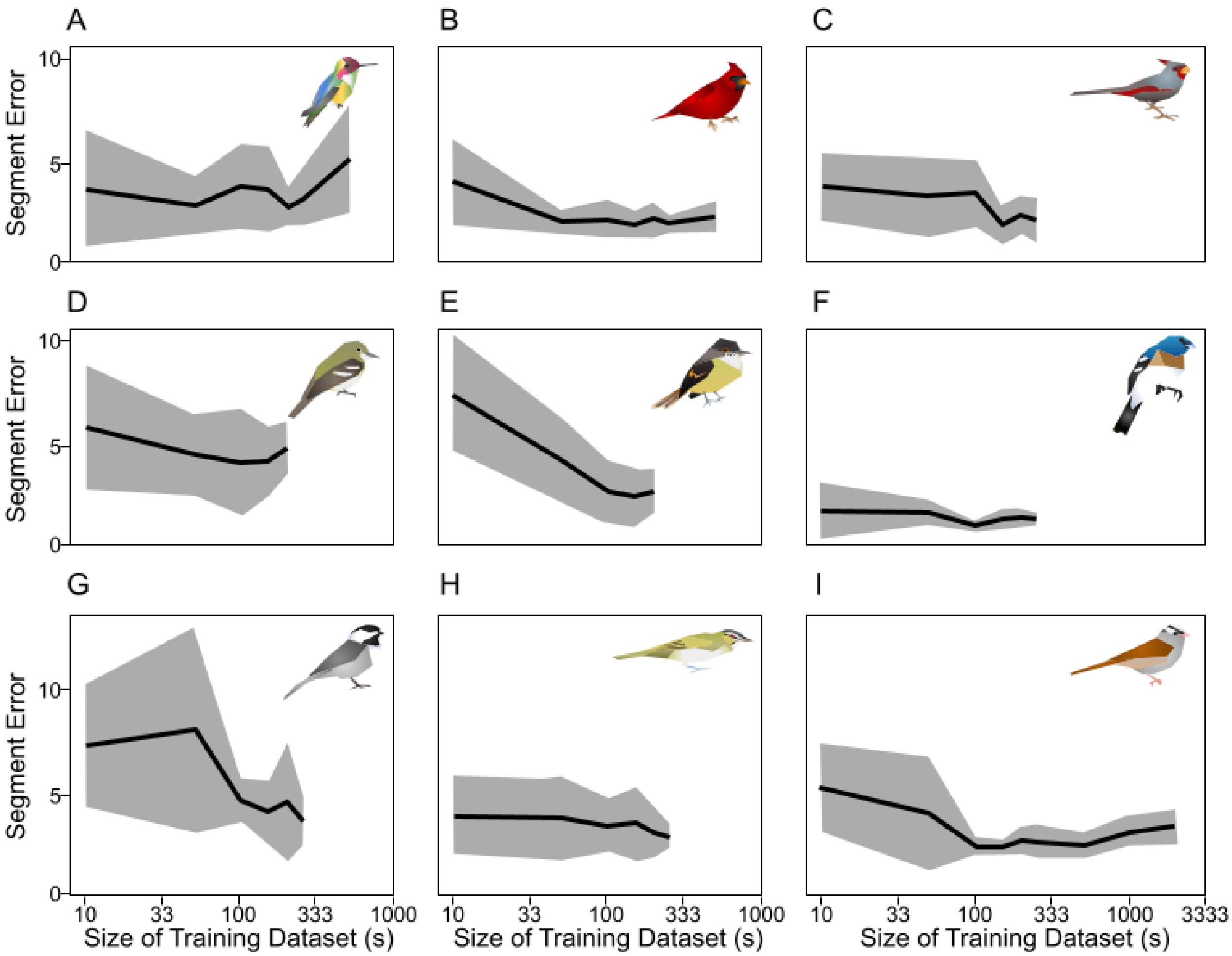

**S3 Fig: Segment error rate of models improves with larger training data, but plateaus quickly**. X-axis gives the size of the training dataset in seconds (log scale). Y-axis gives the uncorrected segment error rate, with higher values being worse performance. Solid line connects mean estimates across 10 replicates of data, with gray polygons giving one standard deviation. A) *Calypte anna*. B) *Cardinalis cardinalis*. C) *Cardinalis sinuatus*. D) *Empidonax virescens*. E) *Myiarchus tuberculifer*. F) *Passerina amoena*. G) *Poecile carolinensis*. H) *Vireo altiloquus*. I) *Zonotrichia leucophrys*.

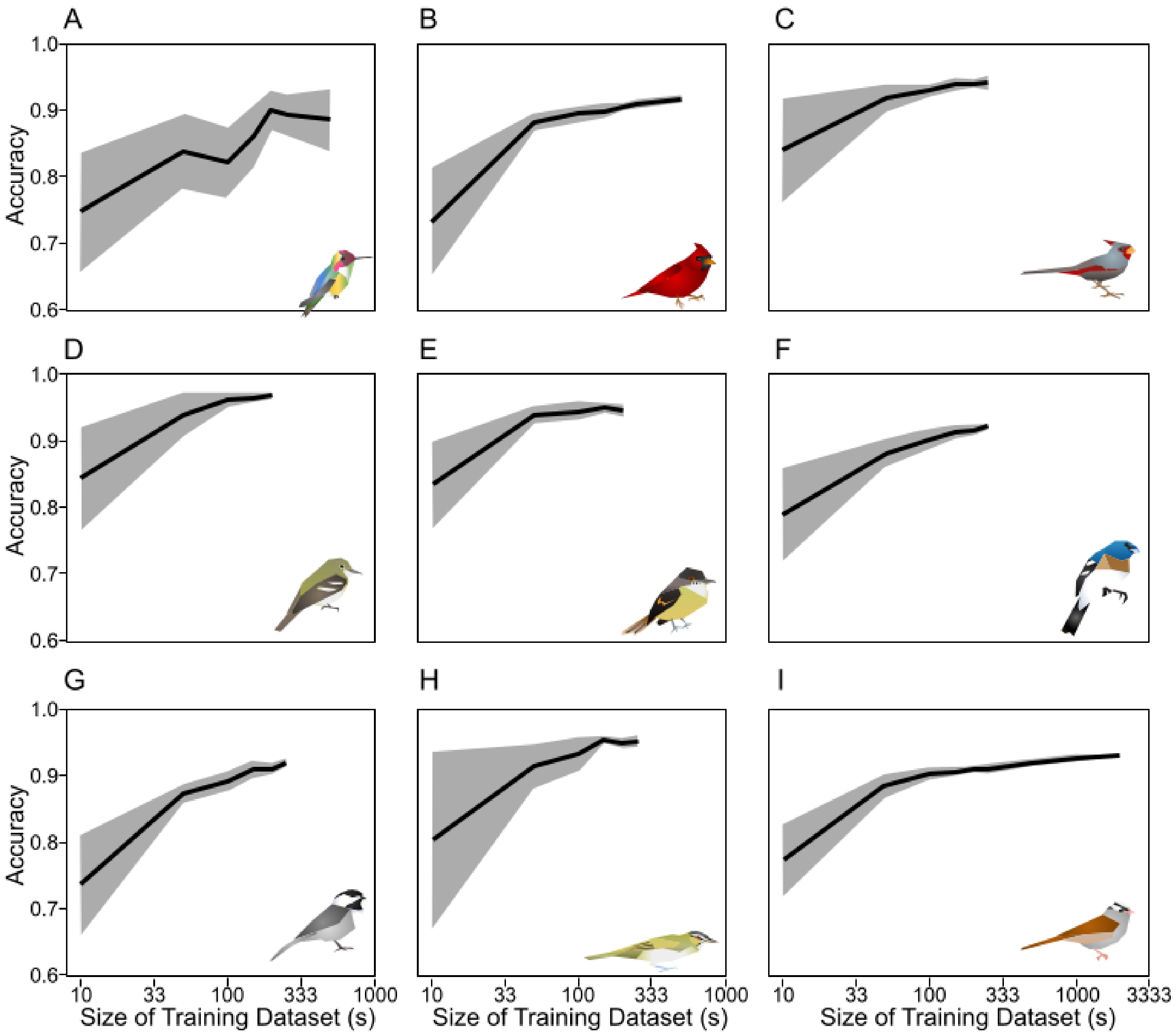

**S4 Fig: Accuracy of models improves with larger training data, but plateaus quickly**. X-axis gives the size of the training dataset in seconds (log scale). Y-axis gives the uncorrected accuracy, with higher values being better performance. Solid line connects mean estimates across 10 replicates of data, with gray polygons giving one standard deviation. A) *Calypte anna*. B) *Cardinalis cardinalis*. C) *Cardinalis sinuatus*. D) *Empidonax virescens*. E) *Myiarchus tuberculifer*. F) *Passerina amoena*. G) *Poecile carolinensis*. H) *Vireo altiloquus*. I) *Zonotrichia leucophrys*.

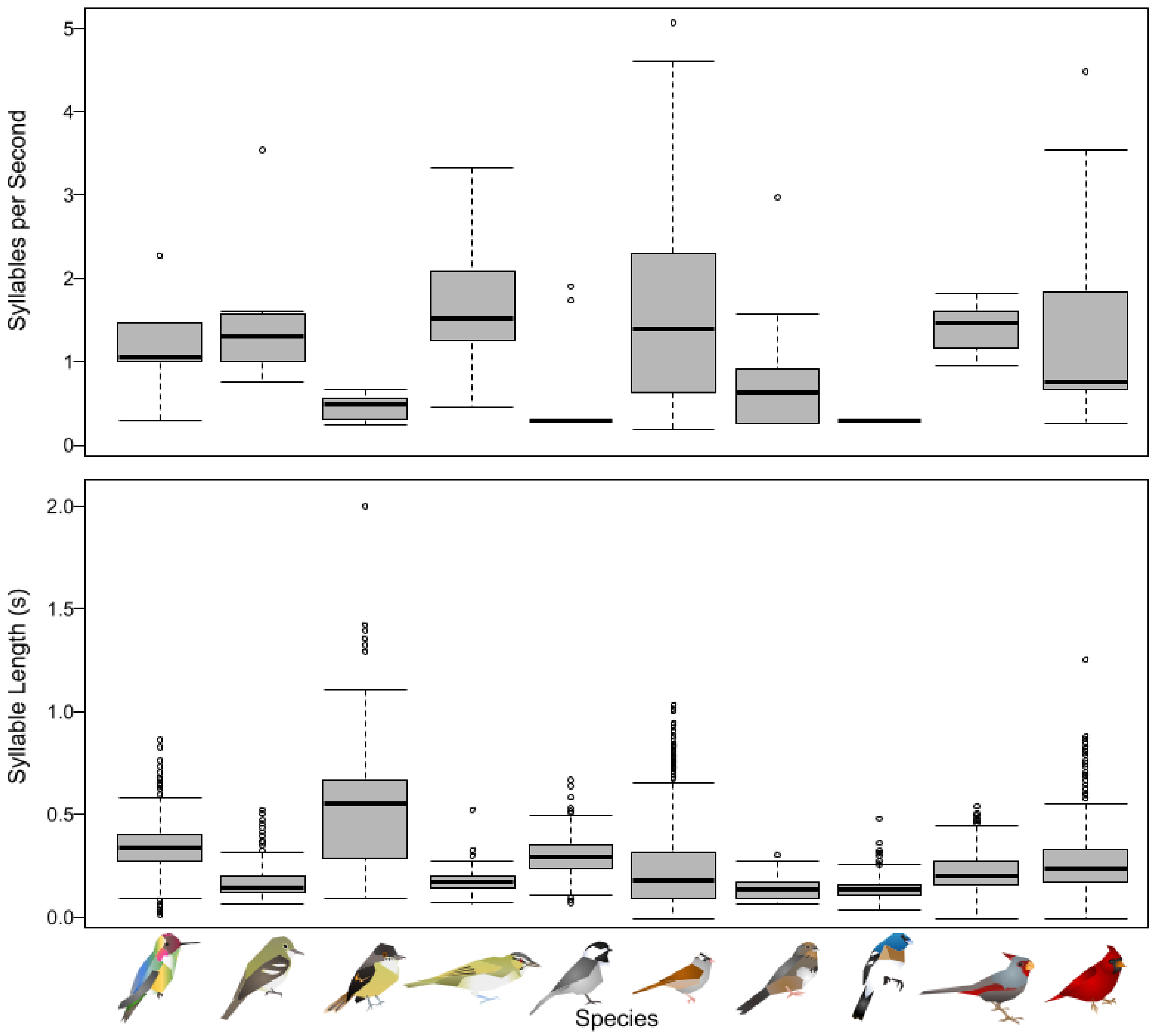

**S4 Fig: Taxa vary in length of syllables as well as number of syllables per second**. X-axis gives species (see Fig 2). Top: boxplot of syllables per second, aka syllable rate. Bottom: boxplot of duration of syllables in seconds.

### Supplementary Tables

**S1 Table: Frame Rate Accuracy and Syllable Error Rate of models trained in TweetyNet across species.**

| Train | Test | Acc | Prec | Rec | F | S.E.R. | Div |
|---|---|---|---|---|---|---|---|
| *9 Species* | *9 Species* | 0.91 (0.91+0.07) | 0.92 (0.90+0.16) | 0.84 (0.85+0.14) | 0.88 (0.86+0.14) | 0.20 (0.64+1.19) | 0 |
| *9 Species* | *C. anna* | 0.78 (0.81+0.13) | 0.93 (0.79+0.36) | 0.66 (0.71+0.21) | 0.78 (0.68+0.29) | 1.44 (2.28+2.09) | 75.6 |
| *9 Species* | *C. cardinalis* | 0.89 (0.90+0.06) | 0.95 (0.95+0.06) | 0.81 (0.77+0.18) | 0.87 (0.84+0.13) | 0.36 (0.49+0.63) | 42.1 |
| *9 Species* | *C. sinuatus* | 0.93 (0.93+0.05) | 0.94 (0.94+0.08) | 0.86 (0.87+0.14) | 0.90 (0.89+0.08) | 0.27 (0.35+0.36) | 42.1 |
| *9 Species* | *E. virescens* | 0.97 (0.97+0.03) | 0.87 (0.90+0.12) | 0.86 (0.88+0.12) | 0.87 (0.88+0.08) | 0.63 (0.79+1.05) | 55.7 |
| *9 Species* | *M. fusca* | 0.94 (0.94+0.05) | 0.87 (0.79+0.26) | 0.91 (0.89+0.09) | 0.89 (0.81+0.20) | 0.19 (1.00+1.46) | 49 |
| *9 Species* | *M. tuberculifer* | 0.94 (0.95+0.03) | 0.92 (0.95+0.07) | 0.91 (0.89+0.09) | 0.92 (0.91+0.05) | 0.31 (0.48+0.47) | 55.8 |
| *9 Species* | *P. amoena* | 0.87 (0.87+0.07) | 0.96 (0.96+0.04) | 0.72 (0.74+0.13) | 0.82 (0.83+0.08) | 0.09 (0.20+0.21) | 42.8 |
| *9 Species* | *P. carolinensis* | 0.93 (0.94+0.05) | 0.90 (0.89+0.15) | 0.90 (0.89+0.13) | 0.90 (0.88+0.11) | 0.41 (0.62+0.80) | 54.8 |
| *9 Species* | *V. altiloquus* | 0.96 (0.96+0.05) | 0.97 (0.97+0.05) | 0.79 (0.82+0.17) | 0.87 (0.88+0.12) | 0.13 (0.18+0.32) | 56.1 |
| *9 Species* | *Z. leucophrys* | 0.90 (0.90+0.08) | 0.93 (0.93+0.11) | 0.84 (0.85+0.15) | 0.88 (0.87+0.12) | 0.13 (0.55+1.23) | 48.1 |
| *C. anna* | *9 Species* | 0.68 (0.69+0.16) | 0.69 (0.76+0.28) | 0.35 (0.39+0.29) | 0.46 (0.44+0.24) | 0.07 (1.13+2.18) | 75.6 |
| *C. anna* | *C. anna* | 0.90 (0.91+0.06) | 0.94 (0.81+0.34) | 0.88 (0.91+0.05) | 0.91 (0.81+0.26) | 0.58 (1.46+2.02) | 0 |
| *C. anna* | *C. cardinalis* | 0.62 (0.65+0.17) | 0.78 (0.87+0.23) | 0.21 (0.27+0.19) | 0.33 (0.38+0.22) | 0.13 (1.12+1.91) | 85 |
| *C. anna* | *C. sinuatus* | 0.65 (0.66+0.08) | 0.64 (0.76+0.32) | 0.15 (0.17+0.18) | 0.25 (0.24+0.21) | 0.16 (0.78+0.85) | 85 |
| *C. anna* | *E. virescens* | 0.81 (0.81+0.23) | 0.33 (0.71+0.35) | 0.52 (0.54+0.36) | 0.40 (0.45+0.23) | 0.89 (1.49+1.93) | 85 |
| *C. anna* | *M. fusca* | 0.80 (0.82+0.15) | 0.65 (0.69+0.32) | 0.37 (0.46+0.31) | 0.47 (0.49+0.28) | 0.18 (1.50+2.93) | 85 |
| *C. anna* | *M. tuberculifer* | 0.56 (0.56+0.21) | 0.37 (0.42+0.24) | 0.35 (0.54+0.35) | 0.36 (0.37+0.22) | 1.86 (5.93+7.11) | 85 |
| *C. anna* | *P. amoena* | 0.69 (0.70+0.10) | 0.91 (0.93+0.06) | 0.31 (0.33+0.24) | 0.46 (0.44+0.23) | 0.30 (0.41+0.21) | 85 |
| *C. anna* | *P. carolinensis* | 0.72 (0.74+0.21) | 0.56 (0.65+0.31) | 0.72 (0.68+0.32) | 0.63 (0.65+0.16) | 0.46 (0.94+0.72) | 85 |
| *C. anna* | *V. altiloquus* | 0.82 (0.82+0.04) | 0.40 (0.30+0.27) | 0.19 (0.18+0.20) | 0.26 (0.33+0.18) | 0.10 (1.12+0.62) | 85 |
| *C. anna* | *Z. leucophrys* | 0.64 (0.64+0.12) | 0.70 (0.80+0.23) | 0.33 (0.35+0.25) | 0.45 (0.42+0.21) | 0.02 (0.76+0.95) | 85 |
| *C. cardinalis* | *9 Species* | 0.85 (0.86+0.12) | 0.87 (0.85+0.21) | 0.73 (0.77+0.23) | 0.79 (0.77+0.20) | 0.26 (0.96+1.56) | 42.1 |
| *C. cardinalis* | *C. anna* | 0.40 (0.48+0.13) | 0.43 (0.35+0.42) | 0.07 (0.09+0.08) | 0.12 (0.08+0.07) | 0.33 (2.19+3.25) | 85 |
| *C. cardinalis* | *C. cardinalis* | 0.92 (0.92+0.05) | 0.95 (0.96+0.05) | 0.86 (0.82+0.18) | 0.90 (0.87+0.13) | 0.31 (0.37+0.49) | 0 |

| | | | | | | | |
|---|---|---|---|---|---|---|---|
| *C. cardinalis* | *C. sinuatus* | 0.92 (0.92+0.06) | 0.92 (0.92+0.09) | 0.87 (0.88+0.18) | 0.89 (0.88+0.12) | 0.11 (0.23+0.19) | 5 |
| *C. cardinalis* | *E. virescens* | 0.90 (0.91+0.09) | 0.57 (0.72+0.28) | 0.91 (0.91+0.10) | 0.70 (0.76+0.19) | 2.11 (1.82+1.76) | 65 |
| *C. cardinalis* | *M. fusca* | 0.92 (0.91+0.05) | 0.77 (0.69+0.27) | 0.95 (0.93+0.08) | 0.85 (0.76+0.21) | 0.10 (1.38+1.85) | 35 |
| *C. cardinalis* | *M. tuberculifer* | 0.87 (0.88+0.07) | 0.81 (0.88+0.18) | 0.83 (0.78+0.21) | 0.82 (0.79+0.12) | 1.90 (2.99+3.23) | 65 |
| *C. cardinalis* | *P. amoena* | 0.76 (0.77+0.12) | 0.95 (0.96+0.05) | 0.48 (0.52+0.22) | 0.63 (0.64+0.19) | 0.12 (0.21+0.25) | 11 |
| *C. cardinalis* | *P. carolinensis* | 0.82 (0.84+0.17) | 0.80 (0.76+0.31) | 0.63 (0.67+0.34) | 0.70 (0.74+0.24) | 0.74 (0.99+0.93) | 55 |
| *C. cardinalis* | *V. altiloquus* | 0.85 (0.86+0.16) | 0.57 (0.73+0.26) | 0.88 (0.89+0.14) | 0.69 (0.77+0.19) | 1.90 (1.62+2.38) | 58 |
| *C. cardinalis* | *Z. leucophrys* | 0.84 (0.84+0.11) | 0.90 (0.91+0.13) | 0.71 (0.71+0.22) | 0.80 (0.77+0.19) | 0.24 (0.71+1.16) | 35 |
| *C. sinuatus* | *9 Species* | 0.83 (0.84+0.13) | 0.83 (0.83+0.20) | 0.71 (0.75+0.24) | 0.77 (0.74+0.20) | 0.24 (0.78+1.26) | 42.1 |
| *C. sinuatus* | *C. anna* | 0.67 (0.75+0.20) | 0.91 (0.71+0.37) | 0.47 (0.64+0.38) | 0.62 (0.57+0.37) | 0.33 (1.24+1.56) | 85 |
| *C. sinuatus* | *C. cardinalis* | 0.88 (0.88+0.09) | 0.91 (0.93+0.08) | 0.81 (0.77+0.25) | 0.86 (0.81+0.21) | 0.22 (0.44+0.70) | 5 |
| *C. sinuatus* | *C. sinuatus* | 0.95 (0.95+0.03) | 0.94 (0.93+0.08) | 0.92 (0.93+0.07) | 0.93 (0.92+0.05) | 0.19 (0.27+0.29) | 0 |
| *C. sinuatus* | *E. virescens* | 0.93 (0.93+0.06) | 0.67 (0.71+0.24) | 0.90 (0.90+0.09) | 0.76 (0.78+0.16) | 1.42 (1.33+1.48) | 65 |
| *C. sinuatus* | *M. fusca* | 0.93 (0.93+0.05) | 0.83 (0.76+0.27) | 0.89 (0.89+0.08) | 0.86 (0.79+0.20) | 0.23 (1.03+1.95) | 35 |
| *C. sinuatus* | *M. tuberculifer* | 0.69 (0.71+0.29) | 0.54 (0.70+0.31) | 0.89 (0.86+0.16) | 0.67 (0.71+0.20) | 0.62 (1.37+1.56) | 65 |
| *C. sinuatus* | *P. amoena* | 0.81 (0.81+0.08) | 0.85 (0.87+0.11) | 0.68 (0.68+0.21) | 0.76 (0.74+0.11) | 0.27 (0.33+0.31) | 11 |
| *C. sinuatus* | *P. carolinensis* | 0.83 (0.84+0.13) | 0.70 (0.74+0.17) | 0.88 (0.88+0.13) | 0.78 (0.79+0.13) | 0.83 (0.95+0.81) | 55 |
| *C. sinuatus* | *V. altiloquus* | 0.83 (0.84+0.16) | 0.53 (0.68+0.30) | 0.81 (0.85+0.18) | 0.64 (0.71+0.21) | 1.47 (1.49+1.65) | 58 |
| *C. sinuatus* | *Z. leucophrys* | 0.79 (0.80+0.12) | 0.85 (0.87+0.14) | 0.65 (0.65+0.25) | 0.74 (0.71+0.21) | 0.21 (0.64+0.95) | 35 |
| *E. virescens* | *9 Species* | 0.77 (0.79+0.13) | 0.92 (0.91+0.18) | 0.45 (0.52+0.27) | 0.60 (0.61+0.23) | 0.25 (0.81+1.25) | 55.7 |
| *E. virescens* | *C. anna* | 0.41 (0.55+0.23) | 0.77 (0.48+0.49) | 0.09 (0.04+0.06) | 0.17 (0.12+0.11) | 0.47 (1.23+1.02) | 85 |
| *E. virescens* | *C. cardinalis* | 0.70 (0.72+0.15) | 0.99 (0.96+0.16) | 0.34 (0.37+0.23) | 0.51 (0.50+0.26) | 0.20 (0.47+0.46) | 65 |
| *E. virescens* | *C. sinuatus* | 0.80 (0.80+0.04) | 0.97 (0.97+0.03) | 0.47 (0.46+0.12) | 0.63 (0.61+0.12) | 0.38 (0.53+0.35) | 65 |
| *E. virescens* | *E. virescens* | 0.97 (0.97+0.03) | 0.97 (0.97+0.05) | 0.79 (0.80+0.16) | 0.87 (0.87+0.11) | 0.63 (0.63+0.71) | 0 |
| *E. virescens* | *M. fusca* | 0.91 (0.91+0.06) | 0.86 (0.80+0.26) | 0.77 (0.79+0.19) | 0.81 (0.75+0.19) | 0.18 (0.96+1.17) | 65 |
| *E. virescens* | *M. tuberculifer* | 0.79 (0.82+0.14) | 0.85 (0.89+0.15) | 0.53 (0.63+0.24) | 0.65 (0.70+0.13) | 1.66 (3.42+3.73) | 27 |
| *E. virescens* | *P. amoena* | 0.73 (0.74+0.13) | 0.92 (0.95+0.07) | 0.39 (0.44+0.22) | 0.55 (0.56+0.26) | 0.15 (0.35+0.27) | 65 |

| | | | | | | | |
|---|---|---|---|---|---|---|---|
| *E. virescens* | *P. carolinensis* | 0.83 (0.84+0.12) | 0.83 (0.87+0.20) | 0.61 (0.60+0.21) | 0.71 (0.69+0.17) | 1.48 (1.49+1.76) | 65 |
| *E. virescens* | *V. altiloquus* | 0.92 (0.92+0.07) | 0.93 (0.93+0.09) | 0.62 (0.65+0.23) | 0.75 (0.74+0.19) | 0.40 (0.60+0.43) | 65 |
| *E. virescens* | *Z. leucophrys* | 0.73 (0.73+0.11) | 0.94 (0.95+0.14) | 0.42 (0.41+0.22) | 0.58 (0.54+0.22) | 0.25 (0.65+0.95) | 65 |
| *9 Species+M. fusca* | *9 Species* | 0.89 (0.90+0.10) | 0.88 (0.90+0.14) | 0.82 (0.84+0.18) | 0.85 (0.85+0.15) | 0.36 (0.73+1.23) | 49 |
| *9 Species+M. fusca* | *C. anna* | 0.74 (0.79+0.16) | 0.91 (0.73+0.23) | 0.61 (0.67+0.33) | 0.73 (0.63+0.28) | 1.81 (2.23+1.32) | 85 |
| *9 Species+M. fusca* | *C. cardinalis* | 0.88 (0.89+0.06) | 0.94 (0.95+0.07) | 0.79 (0.79+0.15) | 0.86 (0.85+0.10) | 0.42 (0.64+1.08) | 35 |
| *9 Species+M. fusca* | *C. sinuatus* | 0.91 (0.91+0.06) | 0.96 (0.96+0.07) | 0.78 (0.79+0.18) | 0.86 (0.85+0.13) | 0.22 (0.34+0.31) | 35 |
| *9 Species+M. fusca* | *E. virescens* | 0.95 (0.95+0.04) | 0.75 (0.81+0.17) | 0.88 (0.88+0.10) | 0.81 (0.82+0.09) | 1.37 (1.25+1.14) | 65 |
| *9 Species+M. fusca* | *M. fusca* | 0.98 (0.98+0.02) | 0.97 (0.96+0.09) | 0.95 (0.93+0.08) | 0.96 (0.94+0.07) | 0.08 (0.18+0.53) | 0 |
| *9 Species+M. fusca* | *M. tuberculifer* | 0.88 (0.90+0.08) | 0.83 (0.91+0.12) | 0.85 (0.80+0.21) | 0.84 (0.83+0.14) | 1.72 (2.73+3.23) | 65 |
| *9 Species+M. fusca* | *P. amoena* | 0.85 (0.84+0.07) | 0.89 (0.90+0.10) | 0.75 (0.73+0.19) | 0.81 (0.79+0.11) | 0.05 (0.35+0.26) | 35 |
| *9 Species+M. fusca* | *P. carolinensis* | 0.91 (0.91+0.05) | 0.88 (0.87+0.16) | 0.86 (0.87+0.13) | 0.87 (0.85+0.11) | 0.72 (0.98+0.89) | 55 |
| *9 Species+M. fusca* | *V. altiloquus* | 0.90 (0.90+0.11) | 0.69 (0.81+0.20) | 0.86 (0.85+0.24) | 0.76 (0.78+0.21) | 0.70 (0.79+0.83) | 58 |
| *9 Species+M. fusca* | *Z. leucophrys* | 0.86 (0.87+0.11) | 0.87 (0.88+0.14) | 0.82 (0.82+0.19) | 0.84 (0.83+0.16) | 0.32 (0.76+1.19) | 8 |
| *M. tuberculifer* | *9 Species* | 0.78 (0.79+0.12) | 0.89 (0.87+0.23) | 0.50 (0.54+0.25) | 0.64 (0.62+0.22) | 0.03 (0.66+0.94) | 55.8 |
| *M. tuberculifer* | *C. anna* | 0.37 (0.49+0.21) | 0.44 (0.49+0.47) | 0.03 (0.02+0.02) | 0.06 (0.06+0.01) | 0.82 (0.92+0.11) | 85 |
| *M. tuberculifer* | *C. cardinalis* | 0.84 (0.85+0.08) | 0.95 (0.95+0.06) | 0.69 (0.71+0.16) | 0.80 (0.80+0.11) | 0.09 (0.40+0.35) | 65 |
| *M. tuberculifer* | *C. sinuatus* | 0.85 (0.85+0.05) | 0.94 (0.95+0.07) | 0.63 (0.66+0.16) | 0.75 (0.76+0.10) | 0.30 (0.28+0.21) | 65 |
| *M. tuberculifer* | *E. virescens* | 0.85 (0.85+0.19) | 0.45 (0.72+0.30) | 0.64 (0.65+0.25) | 0.53 (0.61+0.23) | 1.32 (1.24+0.97) | 27 |
| *M. tuberculifer* | *M. fusca* | 0.88 (0.88+0.06) | 0.82 (0.73+0.32) | 0.66 (0.65+0.20) | 0.73 (0.66+0.22) | 0.22 (1.05+1.52) | 65 |
| *M. tuberculifer* | *M. tuberculifer* | 0.95 (0.95+0.04) | 0.93 (0.96+0.06) | 0.92 (0.92+0.10) | 0.92 (0.93+0.06) | 0.03 (0.18+0.32) | 0 |
| *M. tuberculifer* | *P. amoena* | 0.64 (0.65+0.10) | 0.97 (0.97+0.06) | 0.16 (0.19+0.12) | 0.27 (0.30+0.16) | 0.51 (0.54+0.14) | 65 |
| *M. tuberculifer* | *P. carolinensis* | 0.80 (0.81+0.10) | 0.83 (0.81+0.25) | 0.54 (0.54+0.25) | 0.66 (0.62+0.24) | 0.76 (0.91+0.77) | 65 |
| *M. tuberculifer* | *V. altiloquus* | 0.89 (0.90+0.11) | 0.67 (0.78+0.21) | 0.81 (0.83+0.12) | 0.73 (0.78+0.13) | 0.90 (0.74+0.95) | 65 |
| *M. tuberculifer* | *Z. leucophrys* | 0.74 (0.74+0.10) | 0.93 (0.92+0.18) | 0.45 (0.44+0.21) | 0.60 (0.58+0.20) | 0.02 (0.55+0.74) | 65 |
| *P. amoena* | *9 Species* | 0.85 (0.87+0.12) | 0.87 (0.86+0.20) | 0.73 (0.77+0.21) | 0.80 (0.78+0.20) | 0.25 (0.82+1.48) | 42.8 |
| *P. amoena* | *C. anna* | 0.68 (0.72+0.20) | 0.88 (0.67+0.43) | 0.51 (0.58+0.33) | 0.64 (0.54+0.41) | 0.88 (1.45+1.45) | 85 |

| | | | | | | | |
|---|---|---|---|---|---|---|---|
| *P. amoena* | *C. cardinalis* | 0.68 (0.72+0.16) | 0.87 (0.90+0.16) | 0.35 (0.42+0.21) | 0.50 (0.54+0.23) | 0.12 (0.77+0.81) | 11 |
| *P. amoena* | *C. sinuatus* | 0.79 (0.79+0.05) | 0.92 (0.91+0.11) | 0.47 (0.46+0.13) | 0.62 (0.61+0.13) | 0.36 (0.40+0.33) | 11 |
| *P. amoena* | *E. virescens* | 0.94 (0.94+0.04) | 0.71 (0.75+0.20) | 0.87 (0.87+0.10) | 0.78 (0.79+0.12) | 1.47 (1.72+1.77) | 65 |
| *P. amoena* | *M. fusca* | 0.93 (0.93+0.05) | 0.84 (0.78+0.29) | 0.89 (0.90+0.11) | 0.87 (0.79+0.22) | 0.32 (0.99+1.43) | 35 |
| *P. amoena* | *M. tuberculifer* | 0.64 (0.64+0.27) | 0.51 (0.62+0.27) | 0.60 (0.62+0.31) | 0.55 (0.52+0.26) | 2.66 (4.35+4.22) | 65 |
| *P. amoena* | *P. amoena* | 0.93 (0.93+0.05) | 0.95 (0.94+0.04) | 0.87 (0.88+0.08) | 0.91 (0.91+0.05) | 0.04 (0.12+0.10) | 0 |
| *P. amoena* | *P. carolinensis* | 0.87 (0.88+0.09) | 0.77 (0.75+0.18) | 0.88 (0.85+0.22) | 0.82 (0.79+0.19) | 0.83 (0.89+0.68) | 55 |
| *P. amoena* | *V. altiloquus* | 0.90 (0.90+0.06) | 0.85 (0.91+0.15) | 0.52 (0.53+0.23) | 0.65 (0.64+0.21) | 0.57 (0.56+0.70) | 58 |
| *P. amoena* | *Z. leucophrys* | 0.87 (0.87+0.09) | 0.90 (0.91+0.12) | 0.79 (0.79+0.16) | 0.84 (0.83+0.12) | 0.14 (0.56+1.09) | 35 |
| *P. carolinensis* | *9 Species* | 0.80 (0.82+0.12) | 0.95 (0.93+0.16) | 0.52 (0.55+0.22) | 0.67 (0.66+0.20) | 0.10 (0.61+1.25) | 54.8 |
| *P. carolinensis* | *C. anna* | 0.50 (0.60+0.16) | 0.80 (0.68+0.45) | 0.14 (0.16+0.14) | 0.24 (0.19+0.19) | 0.16 (1.13+1.65) | 85 |
| *P. carolinensis* | *C. cardinalis* | 0.70 (0.72+0.15) | 0.97 (0.96+0.13) | 0.35 (0.37+0.25) | 0.51 (0.50+0.27) | 0.08 (0.75+0.61) | 55 |
| *P. carolinensis* | *C. sinuatus* | 0.77 (0.77+0.04) | 0.97 (0.96+0.07) | 0.40 (0.39+0.10) | 0.56 (0.55+0.11) | 0.20 (0.31+0.30) | 55 |
| *P. carolinensis* | *E. virescens* | 0.90 (0.90+0.14) | 0.62 (0.87+0.25) | 0.61 (0.61+0.23) | 0.61 (0.66+0.20) | 1.37 (1.22+2.22) | 65 |
| *P. carolinensis* | *M. fusca* | 0.91 (0.92+0.06) | 0.96 (0.88+0.23) | 0.68 (0.62+0.23) | 0.80 (0.71+0.20) | 0.24 (0.59+0.74) | 55 |
| *P. carolinensis* | *M. tuberculifer* | 0.80 (0.83+0.13) | 0.86 (0.83+0.18) | 0.53 (0.60+0.23) | 0.66 (0.68+0.18) | 1.38 (2.21+2.42) | 65 |
| *P. carolinensis* | *P. amoena* | 0.75 (0.76+0.11) | 0.94 (0.95+0.06) | 0.44 (0.48+0.20) | 0.60 (0.61+0.16) | 0.18 (0.33+0.29) | 55 |
| *P. carolinensis* | *P. carolinensis* | 0.95 (0.95+0.04) | 0.95 (0.93+0.14) | 0.90 (0.88+0.12) | 0.92 (0.89+0.11) | 0.17 (0.30+0.57) | 0 |
| *P. carolinensis* | *V. altiloquus* | 0.88 (0.88+0.07) | 0.78 (0.90+0.18) | 0.50 (0.49+0.25) | 0.61 (0.59+0.22) | 0.63 (0.83+0.81) | 58 |
| *P. carolinensis* | *Z. leucophrys* | 0.78 (0.79+0.10) | 0.96 (0.96+0.10) | 0.54 (0.55+0.18) | 0.69 (0.68+0.16) | 0.04 (0.49+1.32) | 55 |
| *V. altiloquus* | *9 Species* | 0.83 (0.84+0.13) | 0.86 (0.85+0.24) | 0.70 (0.73+0.25) | 0.77 (0.75+0.21) | 0.27 (1.05+1.98) | 56.1 |
| *V. altiloquus* | *C. anna* | 0.37 (0.41+0.13) | 0.44 (0.30+0.47) | 0.10 (0.28+0.40) | 0.17 (0.15+0.06) | 0.02 (2.45+4.25) | 85 |
| *V. altiloquus* | *C. cardinalis* | 0.74 (0.76+0.12) | 0.91 (0.92+0.12) | 0.51 (0.55+0.15) | 0.65 (0.67+0.13) | 0.50 (1.02+1.33) | 58 |
| *V. altiloquus* | *C. sinuatus* | 0.80 (0.79+0.07) | 0.87 (0.87+0.19) | 0.54 (0.54+0.14) | 0.67 (0.65+0.14) | 0.33 (0.44+0.65) | 58 |
| *V. altiloquus* | *E. virescens* | 0.90 (0.90+0.20) | 0.59 (0.81+0.23) | 0.81 (0.81+0.20) | 0.69 (0.77+0.21) | 1.42 (1.42+2.37) | 65 |
| *V. altiloquus* | *M. fusca* | 0.87 (0.85+0.15) | 0.69 (0.63+0.38) | 0.86 (0.78+0.29) | 0.77 (0.70+0.29) | 0.55 (2.19+3.32) | 58 |
| *V. altiloquus* | *M. tuberculifer* | 0.90 (0.91+0.05) | 0.86 (0.87+0.10) | 0.88 (0.87+0.13) | 0.87 (0.86+0.08) | 1.21 (2.03+2.02) | 65 |

| | | | | | | | |
|---|---|---|---|---|---|---|---|
| *V. altiloquus* | *P. amoena* | 0.79 (0.80+0.08) | 0.88 (0.89+0.10) | 0.60 (0.62+0.19) | 0.71 (0.71+0.15) | 0.00 (0.22+0.16) | 58 |
| *V. altiloquus* | *P. carolinensis* | 0.82 (0.84+0.13) | 0.70 (0.78+0.20) | 0.82 (0.83+0.12) | 0.76 (0.78+0.11) | 1.46 (1.55+1.06) | 58 |
| *V. altiloquus* | *V. altiloquus* | 0.95 (0.95+0.04) | 0.89 (0.93+0.09) | 0.84 (0.85+0.15) | 0.87 (0.87+0.10) | 0.23 (0.22+0.33) | 0 |
| *V. altiloquus* | *Z. leucophrys* | 0.85 (0.85+0.12) | 0.92 (0.93+0.12) | 0.73 (0.72+0.24) | 0.81 (0.78+0.20) | 0.12 (0.63+1.18) | 58 |
| *Z. leucophrys* | *9 Species* | 0.88 (0.90+0.11) | 0.91 (0.89+0.20) | 0.78 (0.78+0.24) | 0.84 (0.80+0.21) | 0.07 (0.73+1.97) | 48.1 |
| *Z. leucophrys* | *C. anna* | 0.39 (0.46+0.11) | 0.58 (0.43+0.49) | 0.10 (0.05+0.06) | 0.17 (0.12+0.11) | 0.33 (1.80+2.37) | 85 |
| *Z. leucophrys* | *C. cardinalis* | 0.69 (0.71+0.13) | 0.94 (0.94+0.10) | 0.35 (0.39+0.19) | 0.51 (0.53+0.20) | 0.18 (0.67+0.54) | 35 |
| *Z. leucophrys* | *C. sinuatus* | 0.81 (0.81+0.06) | 0.93 (0.93+0.13) | 0.53 (0.52+0.13) | 0.67 (0.65+0.12) | 0.25 (0.40+0.36) | 35 |
| *Z. leucophrys* | *E. virescens* | 0.91 (0.92+0.06) | 0.63 (0.77+0.26) | 0.81 (0.80+0.25) | 0.71 (0.72+0.22) | 0.89 (1.00+0.80) | 65 |
| *Z. leucophrys* | *M. fusca* | 0.92 (0.92+0.05) | 0.86 (0.77+0.32) | 0.82 (0.68+0.24) | 0.84 (0.69+0.28) | 0.28 (0.98+1.65) | 8 |
| *Z. leucophrys* | *M. tuberculifer* | 0.76 (0.76+0.18) | 0.68 (0.78+0.29) | 0.64 (0.67+0.19) | 0.66 (0.66+0.14) | 2.21 (5.24+9.14) | 65 |
| *Z. leucophrys* | *P. amoena* | 0.83 (0.83+0.07) | 0.91 (0.92+0.08) | 0.66 (0.68+0.16) | 0.77 (0.77+0.09) | 0.19 (0.25+0.20) | 35 |
| *Z. leucophrys* | *P. carolinensis* | 0.82 (0.83+0.15) | 0.71 (0.78+0.21) | 0.80 (0.79+0.11) | 0.75 (0.77+0.12) | 1.48 (1.47+1.53) | 55 |
| *Z. leucophrys* | *V. altiloquus* | 0.93 (0.93+0.06) | 0.81 (0.89+0.15) | 0.78 (0.76+0.23) | 0.79 (0.79+0.16) | 0.60 (0.53+0.67) | 58 |
| *Z. leucophrys* | *Z. leucophrys* | 0.94 (0.94+0.05) | 0.95 (0.95+0.08) | 0.91 (0.91+0.13) | 0.93 (0.92+0.10) | 0.08 (0.34+0.43) | 0 |

Numerical estimates for performance (Acc, Prec, Rec, F, S.E.R) are given as point estimate (mean+standard deviation). “Train” = species model was trained on. “Test” = species model was tested on. “Acc” = model accuracy. “Prec” = model precision. “Rec” = model recall. “F” = model F-score. “S.E.R” = absolute model segment error rate. “Div” = divergence time between trained and tested species. Estimated divergence time for “9 Species” models is a weighted average, proportional to the sample sizes used for each species trained on.

**S2 Table: Statistical significance and model parameters of ANOVA tests.**

| Response | Predictor | F | p | Sum squares | Mean squares |
|---|---|---|---|---|---|
| accuracy | TEST | 10.74 | **1.57x10$^{-12}$** | 1.00 | 0.10 |
| accuracy | TRAIN | 1.92 | *0.049* | 0.30 | 0.030 |
| accuracy (mean) | TEST | 8.11 | **1.03x10$^{-9}$** | 0.66 | 0.066 |
| accuracy (mean) | TRAIN | 2.37 | *0.013* | 0.27 | 0.027 |
| F-score | TEST | 3.42 | *0.00061* | 1.10 | 0.11 |
| F-score | TRAIN | 5.05 | **4.84x10$^{-6}$** | 1.46 | 0.14 |
| F-score (mean) | TEST | 4.81 | **9.84x10$^{-6}$** | 1.36 | 0.13 |
| F-score (mean) | TRAIN | 4.57 | **1.98x10$^{-5}$** | 1.31 | 0.13 |
| precision | TEST | 5.59 | **1.03x10$^{-6}$** | 0.97 | 0.097 |
| precision | TRAIN | 3.62 | **0.00033** | 0.71 | 0.071 |
| precision (mean) | TEST | 9.81 | **1.41x10$^{-11}$** | 1.14 | 0.11 |
| precision (mean) | TRAIN | 2.14 | *0.026* | 0.39 | 0.039 |
| recall | TEST | 4.79 | **1.04x10$^{-5}$** | 2.01 | 0.20 |
| recall | TRAIN | 5.39 | **1.80x10$^{-6}$** | 2.18 | 0.21 |
| recall (mean) | TEST | 4.18 | **6.30x10$^{-5}$** | 1.64 | 0.16 |
| recall (mean) | TRAIN | 5.44 | **1.59x10$^{-6}$** | 1.97 | 0.19 |
| S.E.R. | TEST | 14.2 | **7.62x10$^{-16}$** | 225828 | 22583 |
| S.E.R. | TRAIN | 0.48 | 0.89 | 16952 | 1695 |
| S.E.R. (mean) | TEST | 13.3 | **5.10x10$^{-15}$** | 551384 | 55138 |
| S.E.R. (mean) | TRAIN | 0.85 | 0.57 | 72857 | 7286 |
| Uncorrected accuracy | TEST | 5.52 | **1.25x10$^{-6}$** | 0.41 | 0.041 |
| Uncorrected accuracy | TRAIN | 2.92 | *0.0027* | 0.26 | 0.026 |
| Uncorrected S.E.R. | TEST | 13.39 | **4.19x10$^{-15}$** | 4288 | 428.80 |
| Uncorrected S.E.R. | TRAIN | 2.18 | *0.024* | 1292 | 129.18 |

Degrees of freedom for all analyses in this table is 110. “S.E.R” = absolute model segment error rate.
“italics” = near significant ($p < 0.05$)
“bold” = significant ($p < 0.000413$)

**S3 Table: Statistical significance and model parameters of linear models between species' mean performance and song diversity.**

| Response | Predictor | F | Coef. | Err. | T | p | $R^2$ | $aR^2$ |
|---|---|---|---|---|---|---|---|---|
| accuracy | Test Complexity | 2.05 | 0.016 | 0.011 | 1.43 | 0.21 | 0.29 | 0.14 |
| accuracy | Test Hypervolume (mean) | 0.27 | 0.062 | 0.11 | 0.52 | 0.62 | 0.051 | -0.13 |
| accuracy | Test Hypervolume (sd) | 0.13 | 0.043 | 0.11 | 0.36 | 0.72 | 0.026 | -0.16 |
| accuracy | Test InVar | 0.010 | -0.0023 | 0.023 | -0.10 | 0.92 | 0.0020 | -0.19 |
| accuracy | Test SpVar | 2.54 | -0.021 | 0.013 | -1.59 | 0.17 | 0.33 | 0.20 |
| accuracy | Train Complexity | 8.50 | -0.011 | 0.0037 | -2.91 | *0.033* | 0.62 | 0.55 |
| accuracy | Train Hypervolume (mean) | 1.32 | -0.056 | 0.048 | -1.15 | 0.30 | 0.21 | 0.052 |
| accuracy | Train Hypervolume (sd) | 2.97 | -0.073 | 0.042 | -1.72 | 0.14 | 0.37 | 0.24 |
| accuracy | Train InVar | 3.06 | 0.014 | 0.0081 | 1.75 | 0.14 | 0.38 | 0.25 |
| accuracy | Train SpVar | 2.07 | 0.0090 | 0.0062 | 1.44 | 0.20 | 0.29 | 0.15 |
| F-score | Test Complexity | 0.60 | 0.010 | 0.013 | 0.78 | 0.47 | 0.10 | -0.069 |
| F-score | Test Hypervolume (mean) | 12.94 | 0.23 | 0.064 | 3.59 | *0.015* | 0.72 | 0.66 |
| F-score | Test Hypervolume (sd) | 24.77 | 0.24 | 0.048 | 4.97 | *0.0041* | 0.83 | 0.79 |
| F-score | Test InVar | 2.15 | -0.028 | 0.019 | -1.46 | 0.20 | 0.30 | 0.16 |
| F-score | Test SpVar | 0.69 | -0.012 | 0.015 | -0.83 | 0.44 | 0.12 | -0.053 |
| F-score | Train Complexity | 3.67 | -0.027 | 0.014 | -1.91 | 0.11 | 0.42 | 0.30 |
| F-score | Train Hypervolume (mean) | 2.88 | -0.22 | 0.13 | -1.69 | 0.15 | 0.36 | 0.23 |
| F-score | Train Hypervolume (sd) | 5.60 | -0.26 | 0.11 | -2.36 | 0.064 | 0.52 | 0.43 |
| F-score | Train InVar | 5.15 | 0.050 | 0.022 | 2.27 | 0.072 | 0.50 | 0.40 |
| F-score | Train SpVar | 1.38 | 0.023 | 0.020 | 1.17 | 0.29 | 0.21 | 0.060 |
| precision | Test Complexity | 29.57 | -0.061 | 0.011 | -5.43 | *0.0028* | 0.85 | 0.82 |
| precision | Test Hypervolume (mean) | 0.47 | -0.17 | 0.24 | -0.69 | 0.52 | 0.087 | -0.095 |
| precision | Test Hypervolume (sd) | 0.78 | -0.21 | 0.23 | -0.88 | 0.417 | 0.13 | -0.037 |
| precision | Test InVar | 0.87 | 0.042 | 0.045 | 0.93 | 0.39 | 0.14 | -0.021 |
| precision | Test SpVar | 4.81 | 0.055 | 0.025 | 2.19 | 0.079 | 0.49 | 0.38 |
| precision | Train Complexity | 0.46 | 0.0087 | 0.012 | 0.68 | 0.52 | 0.085 | -0.097 |
| precision | Train Hypervolume (mean) | 28.72 | 0.24 | 0.045 | 5.36 | *0.0030* | 0.85 | 0.82 |
| precision | Train Hypervolume (sd) | 16.57 | 0.22 | 0.055 | 4.07 | *0.0096* | 0.76 | 0.72 |
| precision | Train InVar | 8.85 | -0.039 | 0.013 | -2.97 | *0.031* | 0.63 | 0.56 |
| precision | Train SpVar | 2.28 | -0.019 | 0.013 | -1.51 | 0.19 | 0.31 | 0.17 |
| recall | Test Complexity | 5.83 | 0.048 | 0.020 | 2.41 | 0.060 | 0.53 | 0.44 |
| recall | Test Hypervolume (mean) | 5.15 | 0.41 | 0.18 | 2.27 | 0.072 | 0.50 | 0.40 |
| recall | Test Hypervolume (sd) | 7.65 | 0.44 | 0.16 | 2.76 | *0.039* | 0.60 | 0.52 |
| recall | Test InVar | 2.46 | -0.063 | 0.040 | -1.57 | 0.17 | 0.33 | 0.19 |
| recall | Test SpVar | 3.28 | -0.049 | 0.027 | -1.81 | 0.13 | 0.39 | 0.27 |
| recall | Train Complexity | 3.88 | -0.047 | 0.024 | -1.97 | 0.10 | 0.43 | 0.32 |
| recall | Train Hypervolume (mean) | 6.20 | -0.47 | 0.19 | -2.49 | 0.055 | 0.55 | 0.46 |
| recall | Train Hypervolume (sd) | 11.12 | -0.52 | 0.15 | -3.33 | *0.020* | 0.68 | 0.62 |
| recall | Train InVar | 12.20 | 0.10 | 0.029 | 3.49 | *0.017* | 0.70 | 0.65 |
| recall | Train SpVar | 2.87 | 0.052 | 0.030 | 1.69 | 0.15 | 0.36 | 0.23 |
| S.E.R. | Test Complexity | 8.79 | 29.00 | 9.78 | 2.96 | *0.031* | 0.63 | 0.56 |
| S.E.R. | Test Hypervolume (mean) | 0.36 | 83.39 | 138.64 | 0.60 | 0.57 | 0.067 | -0.11 |
| S.E.R. | Test Hypervolume (sd) | 0.77 | 114.84 | 130.73 | 0.87 | 0.42 | 0.13 | -0.039 |
| S.E.R. | Test InVar | 0.54 | -18.89 | 25.70 | -0.73 | 0.49 | 0.097 | -0.08 |
| S.E.R. | Test SpVar | 1.58 | -21.29 | 16.89 | -1.26 | 0.26 | 0.24 | 0.089 |
| S.E.R. | Train Complexity | 4.02 | -6.01 | 2.99 | -2.00 | 0.10 | 0.44 | 0.33 |
| S.E.R. | Train Hypervolume (mean) | 3.01 | -48.77 | 28.09 | -1.73 | 0.14 | 0.37 | 0.25 |
| S.E.R. | Train Hypervolume (sd) | 4.05 | -52.05 | 25.86 | -2.01 | 0.10 | 0.44 | 0.33 |
| S.E.R. | Train InVar | 0.60 | 4.91 | 6.33 | 0.77 | 0.47 | 0.10 | -0.070 |
| S.E.R. | Train SpVar | 2.29 | 6.02 | 3.97 | 1.51 | 0.19 | 0.31 | 0.17 |

Degrees of freedom for all analyses in this table is 5. “S.E.R” = absolute model segment error rate. “Sd” = standard deviation. “InVar” = individual variation. “SpVar” = species variation. Note that Complexity, InVar, and SpVar are taken from Medina and Francis 2012 [Ref 68]. “Coef” = coefficient value. “Err” = standard error of coefficient. “a$R^2$” = adjusted $R^2$ value.

“italics” = near significant ($p < 0.05$)

“bold” = significant ($p < 0.000413$)

**S4 Table: Calculated hypervolumes correlate highly with pseudo hypervolumes at high levels of dimensionality.**

| **Dimensions** | **F** | **Coef.** | **Err.** | **T** | **p** | **$R^2$** | **$aR^2$** |
|---|---|---|---|---|---|---|---|
| 2 | 0.17 | $1.68 \times 10^{-3}$ | 0.0040 | 0.41 | 0.68 | 0.0035 | -0.017 |
| 3 | 4.24 | $2.32 \times 10^{-3}$ | 0.0011 | 2.06 | *0.044* | 0.081 | 0.062 |
| 4 | 13.67 | $8.98 \times 10^{-4}$ | 0.00024 | 3.69 | *0.00055* | 0.22 | 0.20 |
| 5 | 30.73 | $3.41 \times 10^{-4}$ | 0.000061 | 5.54 | **$1.24 \times 10^{-6}$** | 0.39 | 0.37 |
| 6 | 57.95 | $1.29 \times 10^{-4}$ | 0.000016 | 7.61 | **$8.52 \times 10^{-10}$** | 0.54 | 0.53 |
| 7 | 72.88 | $4.99 \times 10^{-5}$ | 0.0000058 | 8.53 | **$3.44 \times 10^{-11}$** | 0.60 | 0.59 |

Degrees of freedom for all analyses in this table is 48. "Coef" = coefficient value. "Err" = standard error of coefficient. "$aR^2$" = adjusted $R^2$ value.
"italics" = near significant ($p < 0.05$)
"bold" = significant ($p < 0.000413$)

**S5 Table: Statistical significance and model parameters of linear models between non-mean performance values and song diversity, as well as across song-diversity measures.**

| **Response** | **Predictor** | **F** | **Coef.** | **Err.** | **T** | **p** | **$R^2$** | **$aR^2$** |
|---|---|---|---|---|---|---|---|---|
| accuracy | Train Complexity | 0.77 | -0.011 | 0.012 | -0.88 | 0.38 | 0.010 | -0.0029 |
| accuracy | Train InVar | 0.46 | 0.014 | 0.020 | 0.68 | 0.49 | 0.0061 | -0.0070 |
| accuracy | Train SpVar | 0.35 | 0.0090 | 0.015 | 0.59 | 0.55 | 0.0047 | -0.0085 |
| F-score | Train Complexity | 2.51 | -0.027 | 0.017 | -1.58 | 0.11 | 0.032 | 0.019 |
| F-score | Train InVar | 3.03 | 0.050 | 0.028 | 1.74 | 0.085 | 0.038 | 0.026 |
| F-score | Train SpVar | 1.27 | 0.023 | 0.020 | 1.12 | 0.26 | 0.016 | 0.0035 |
| precision | Train Complexity | 0.37 | 0.0087 | 0.014 | 0.61 | 0.54 | 0.0049 | -0.0082 |
| precision | Train InVar | 2.90 | -0.039 | 0.023 | -1.70 | 0.092 | 0.037 | 0.024 |
| precision | Train SpVar | 1.39 | -0.019 | 0.016 | -1.18 | 0.24 | 0.018 | 0.0052 |
| recall | Train Complexity | 5.18 | -0.047 | 0.021 | -2.27 | *0.025* | 0.064 | 0.052 |
| recall | Train InVar | 8.79 | 0.10 | 0.034 | 2.96 | *0.0040* | 0.10 | 0.092 |
| recall | Train SpVar | 4.28 | 0.052 | 0.025 | 2.07 | *0.041* | 0.054 | 0.041 |
| S.E.R. | Train Complexity | 1.10 | -6.01 | 5.71 | -1.05 | 0.29 | 0.014 | 0.0013 |
| S.E.R. | Train InVar | 0.26 | 4.91 | 9.57 | 0.51 | 0.60 | 0.0035 | -0.0097 |
| S.E.R. | Train SpVar | 0.77 | 6.02 | 6.83 | 0.88 | 0.38 | 0.010 | -0.0029 |
| Train Complexity | Train Hypervolume (mean) | 4.48 | 2.09 | 0.99 | 2.11 | *0.037* | 0.056 | 0.043 |
| Train Complexity | Train Hypervolume (sd) | 8.49 | 2.75 | 0.94 | 2.91 | *0.0046* | 0.10 | 0.089 |
| Train Complexity | Train InVar | 19.88 | -0.76 | 0.17 | -4.45 | **$2.84 \times 10^{-5}$** | 0.20 | 0.19 |
| Train Complexity | Train SpVar | 238.80 | -1.04 | 0.067 | -15.45 | **$2.00 \times 10^{-16}$** | 0.76 | 0.75 |
| Train InVar | Train Hypervolume (mean) | 97.84 | -3.99 | 0.40 | -9.89 | **$3.06 \times 10^{-15}$** | 0.56 | 0.56 |
| Train InVar | Train Hypervolume (sd) | 138.50 | -4.18 | 0.35 | -11.76 | **$2.00 \times 10^{-16}$** | 0.64 | 0.64 |
| Train InVar | Train SpVar | 40.85 | 0.42 | 0.066 | 6.39 | **$1.25 \times 10^{-8}$** | 0.35 | 0.34 |
| Train SpVar | Train Hypervolume (mean) | 17.05 | -3.18 | 0.77 | -4.12 | **$9.34 \times 10^{-5}$** | 0.18 | 0.17 |
| Train SpVar | Train Hypervolume (sd) | 16.88 | -3.10 | 0.75 | -4.10 | **0.00010** | 0.18 | 0.17 |
| Training.Time | Train Complexity | 48.94 | -6436 | 920 | -6.99 | **$9.46 \times 10^{-10}$** | 0.39 | 0.38 |
| Training.Time | Train InVar | 0.44 | -1317 | 1964 | -0.67 | 0.505 | 0.0059 | -0.0072 |
| Training.Time | Train SpVar | 70.32 | 8505 | 1014 | 8.38 | **$2.20 \times 10^{-12}$** | 0.48 | 0.47 |

Degrees of freedom for all analyses in this table is 75. “S.E.R” = absolute model segment error rate. “Sd” = standard deviation. “InVar” = individual variation. “SpVar” = species variation. Note that Complexity, InVar, and SpVar are taken from Medina and Francis 2012 [Ref 68]. “Coef” = coefficient value. “Err” = standard error of coefficient. “$aR^2$” = adjusted $R^2$ value.

“italics” = near significant ($p < 0.05$)

“bold” = significant ($p < 0.000413$)

**S6 Table: Statistical significance and model parameters of linear models between non-mean performance values and hypervolume metrics.**

| Response | Predictor | F | Coef. | Err. | T | p | $R^2$ | $aR^2$ |
|---|---|---|---|---|---|---|---|---|
| accuracy | Train Hypervolume (mean) | 6.09 | -0.11 | 0.046 | -2.46 | *0.015* | 0.053 | 0.044 |
| accuracy | Train Hypervolume (sd) | 2.85 | -0.16 | 0.095 | -1.68 | 0.094 | 0.025 | 0.016 |
| F-score | Train Hypervolume (mean) | 21.24 | -0.30 | 0.065 | -4.60 | **$1.11 x 10^{-5}$** | 0.16 | 0.15 |
| F-score | Train Hypervolume (sd) | 10.37 | -0.44 | 0.13 | -3.22 | *0.0016* | 0.08764 | 0.079 |
| precision | Train Hypervolume (mean) | 8.76 | -0.16 | 0.055 | -2.96 | *0.0037* | 0.075 | 0.066 |
| precision | Train Hypervolume (sd) | 0.029 | -0.020 | 0.11 | -0.17 | 0.86 | 0.00027 | -0.0089 |
| recall | Train Hypervolume (mean) | 19.7 | -0.35 | 0.079 | -4.43 | **$2.20 x 10^{-5}$** | 0.15 | 0.14 |
| recall | Train Hypervolume (sd) | 15.22 | -0.64 | 0.16 | -3.90 | **0.00016** | 0.12 | 0.11 |
| S.E.R. | Train Hypervolume (mean) | 1.47 | -25.88 | 21.35 | -1.21 | 0.22 | 0.013 | 0.0042 |
| S.E.R. | Train Hypervolume (sd) | 1.91 | -59.86 | 43.22 | -1.38 | 0.16 | 0.017 | 0.0083 |
| Train Hypervolume (mean) | Train Hypervolume (sd) | 95.89 | 1.39 | 0.14 | 9.79 | *$2.00 x 10^{-16}$* | 0.47 | 0.46 |
| Training.Time | Train Hypervolume (mean) | 33.77 | 24080 | 4144 | 5.81 | *$6.36 x 10^{-8}$* | 0.23 | 0.23 |
| Training.Time | Train Hypervolume (sd) | 3.52 | 17787 | 9478 | 1.87 | 0.063 | 0.031 | 0.022 |

Degrees of freedom for all analyses in this table is 108. “S.E.R” = absolute model segment error rate. “Sd” = standard deviation. “Coef” = coefficient value. “Err” = standard error of coefficient. “$aR^2$” = adjusted $R^2$ value.
“italics” = near significant ($p < 0.05$)
“bold” = significant ($p < 0.000413$)

**S7 Table: Statistical significance and model parameters of linear models between performance metrics, divergence time, training time, and sample size.**

| Response | Predictor | F | Coef. | Err. | T | p | $R^2$ | $aR^2$ |
|---|---|---|---|---|---|---|---|---|
| accuracy | accuracy (mean) | 6054 | 1.12 | 0.014 | 77.81 | **$2.00 \times 10^{-16}$** | 0.98 | 0.98 |
| accuracy | F-score | 465.40 | 0.58 | 0.027 | 21.57 | **$2.00 \times 10^{-16}$** | 0.79 | 0.79 |
| accuracy | precision | 21.31 | 0.32 | 0.070 | 4.616 | **$9.95 \times 10^{-6}$** | 0.15 | 0.14 |
| accuracy | recall | 348.40 | 0.47 | 0.025 | 18.67 | **$2.00 \times 10^{-16}$** | 0.74 | 0.74 |
| accuracy | Testing Test Size | 0.054 | -0.0052 | 0.022 | -0.23 | 0.816 | 0.00045 | -0.0079 |
| accuracy | Testing Train Size | 6.36 | -0.063 | 0.025 | -2.52 | *0.013* | 0.050 | 0.042 |
| accuracy | Testing Val Size | 10.51 | -0.10 | 0.031 | -3.24 | *0.0015* | 0.081 | 0.073 |
| accuracy | Training Test Size | 6.74 | 0.057 | 0.022 | 2.59 | *0.010* | 0.053 | 0.045 |
| accuracy | Training Train Size | 0.049 | 0.0057 | 0.025 | 0.22 | 0.82 | 0.00041 | -0.0079 |
| accuracy | Training Val Size | 0.21 | 0.015 | 0.033 | 0.46 | 0.64 | 0.0017 | -0.0066 |
| accuracy | Uncorrected accuracy | 1698 | 1.22 | 0.029 | 41.20 | **$2.00 \times 10^{-16}$** | 0.93 | 0.93 |
| F-score | F-score (mean) | 2806 | 0.99 | 0.018 | 52.97 | **$2.00 \times 10^{-16}$** | 0.95 | 0.95 |
| F-score | precision | 49.04 | 0.68 | 0.097 | 7.00 | **$1.61 \times 10^{-10}$** | 0.29 | 0.28 |
| F-score | recall | 742.80 | 0.77 | 0.028 | 27.25 | **$2.00 \times 10^{-16}$** | 0.86 | 0.86 |
| F-score | YEAR | 71.68 | -0.0047 | 0.00056 | -8.46 | **$7.71 \times 10^{-14}$** | 0.37 | 0.37 |
| precision | precision (mean) | 246.60 | 0.89 | 0.057 | 15.7 | **$2.00 \times 10^{-16}$** | 0.67 | 0.67 |
| precision | recall | 6.54 | 0.15 | 0.058 | 2.55 | *0.011* | 0.052 | 0.044 |
| precision | YEAR | 35.65 | -0.0029 | 0.00049 | -5.97 | **$2.51 \times 10^{-8}$** | 0.23 | 0.22 |
| recall | recall (mean) | 3681 | 1.03 | 0.017 | 60.66 | **$2.00 \times 10^{-16}$** | 0.96 | 0.96 |
| recall | YEAR | 44.26 | -0.0048 | 0.00073 | -6.65 | **$9.25 \times 10^{-10}$** | 0.27 | 0.26 |
| S.E.R. | accuracy | 0.014 | 4.92 | 40.76 | 0.12 | 0.90 | 0.00012 | -0.0082 |
| S.E.R. | F-score | 0.070 | -7.15 | 26.88 | -0.26 | 0.79 | 0.00059 | -0.0078 |
| S.E.R. | precision | 33.88 | -175.37 | 30.13 | -5.82 | **$5.07 \times 10^{-8}$** | 0.22 | 0.21 |
| S.E.R. | recall | 3.51 | 41.61 | 22.18 | 1.87 | 0.063 | 0.028 | 0.020 |
| S.E.R. | S.E.R. (mean) | 149.5 | 0.47 | 0.038 | 12.22 | **$2.00 \times 10^{-16}$** | 0.55 | 0.55 |
| S.E.R. | YEAR | 10.24 | 0.64 | 0.20 | 3.20 | *0.0017* | 0.079 | 0.071 |
| S.E.R. (Neg) | YEAR | 4.03 | 0.45 | 0.22 | 2.00 | *0.046* | 0.032 | 0.024 |
| Training Time | Training Train Size | 42.54 | 0.13 | 2.09 | 6.52 | **$1.76 \times 10^{-9}$** | 0.26 | 0.25 |
| Training Time | Training Train Size (log) | 72.79 | 0.00038 | 4475 | 8.53 | **$5.43 \times 10^{-14}$** | 0.37 | 0.37 |
| Uncorrected accuracy | YEAR | 53.35 | -0.0022 | 0.00030 | -7.30 | **$3.45 \times 10^{-11}$** | 0.30 | 0.30 |
| Uncorrected S.E.R. | YEAR | 12.87 | 0.10 | 0.027 | 3.58 | *0.00048* | 0.097 | 0.090 |

Degrees of freedom for all analyses in this table is 119. "S.E.R" = absolute model segment error rate. "S.E.R (Neg)" = model segment error rate including negative values as negative. "Sd" = standard deviation. "Coef" = coefficient value. "Err" = standard error of coefficient. "$aR^2$" = adjusted $R^2$ value. Uncorrected values are taken before removing predicted syllables that are too short.

"italics" = near significant ($p < 0.05$)

"bold" = significant ($p < 0.000413$)

**S8 Table: Species-specific correlations between divergence time and performance of machine learning models.**

| **Response** | **Predictor** | **Species** | **F** | **Coef.** | **Err.** | **T** | **p** | **$R^2$** | **$aR^2$** |
|---|---|---|---|---|---|---|---|---|---|
| accuracy | YEAR | 9SPP | 12.99 | -0.0027 | 0.00075 | -3.60 | *0.0057* | 0.59 | 0.54 |
| S.E.R. | YEAR | 9SPP | 0.74 | -0.16 | 0.18 | -0.86 | 0.41 | 0.076 | -0.026 |
| accuracy | YEAR | CA | 5.56 | -0.0046 | 0.0019 | -2.35 | *0.042* | 0.38 | 0.31 |
| S.E.R. | YEAR | CA | 0.00077 | -0.020 | 0.719 | -0.028 | 0.98 | $8.62x10^{-5}$ | -0.11 |
| accuracy | YEAR | CC | 3.58 | -0.0020 | 0.0010 | -1.89 | 0.091 | 0.28 | 0.21 |
| S.E.R. | YEAR | CC | 0.25 | -0.084 | 0.16 | -0.49 | 0.63 | 0.026 | -0.081 |
| accuracy | YEAR | CS | 0.47 | -0.0022 | 0.00078 | -2.84 | 0.19 | 0.47 | 0.41 |
| S.E.R. | YEAR | CS | 0.51 | 0.071 | 0.10 | 0.71 | 0.49 | 0.053 | -0.051 |
| accuracy | YEAR | EV | 1.28 | -0.00074 | 0.00065 | -1.13 | 0.28 | 0.12 | 0.027 |
| S.E.R. | YEAR | EV | 1.16 | 0.63 | 0.59 | 1.08 | 0.30 | 0.11 | 0.016 |
| accuracy | YEAR | MF | 16.91 | -0.0014 | 0.00035 | -4.11 | *0.0026* | 0.65 | 0.61 |
| S.E.R. | YEAR | MF | 0.35 | 0.097 | 0.16 | 0.59 | 0.56 | 0.038 | -0.068 |
| accuracy | YEAR | MT | 3.55 | -0.0029 | 0.0015 | -1.88 | 0.092 | 0.28 | 0.20 |
| S.E.R. | YEAR | MT | 3.51 | 1.85 | 0.98 | 1.87 | 0.093 | 0.28 | 0.20 |
| accuracy | YEAR | PA | 9.14 | -0.0022 | 0.00073 | -3.02 | *0.014* | 0.50 | 0.44 |
| S.E.R. | YEAR | PA | 1.55 | 0.20 | 0.16 | 1.24 | 0.24 | 0.14 | 0.052 |
| accuracy | YEAR | PC | 12.75 | -0.0024 | 0.00069 | -3.57 | *0.0060* | 0.58 | 0.54 |
| S.E.R. | YEAR | PC | 1.35 | 0.79 | 0.68 | 1.16 | 0.27 | 0.13 | 0.034 |
| accuracy | YEAR | VA | 4.01 | -0.0012 | 0.00062 | -2.004 | *0.076* | 0.30 | 0.23 |
| S.E.R. | YEAR | VA | 0.090 | 0.27 | 0.90 | 0.30 | 0.77 | 0.0099 | -0.10 |
| accuracy | YEAR | ZL | 17.05 | -0.0027 | 0.00067 | -4.12 | *0.0025* | 0.65 | 0.61 |
| S.E.R. | YEAR | ZL | 2.52 | -0.18 | 0.11 | -1.58 | 0.14 | 0.21 | 0.13 |

Degrees of freedom for all analyses in this table is 9. "S.E.R" = absolute model segment error rate. "Coef" = coefficient value. "Err" = standard error of coefficient. "$aR^2$" = adjusted $R^2$ value.
"italics" = near significant ($p < 0.05$)
"bold" = significant ($p < 0.000413$)

**S9 Table: Individual songs used in models.**

| Collection | ID | Genus | Species | Latitude | Longitude | Year | Train (s) | Val (s) | Test (s) |
|---|---|---|---|---|---|---|---|---|---|
| BLB | 28330 | *Calypte* | *anna* | 39.75 | -122.6667 | 1995.00 | 4.45483 | 0 | 4.1875 |
| BLB | 33172 | *Calypte* | *anna* | 34.7667 | -118.3833 | 1980.00 | 6.64037 | 0 | 3.4773 |
| BLB | 33565 | *Calypte* | *anna* | 37.7289 | -122.4948 | 1990.00 | 6.79648 | 0 | 0 |
| BLB | 34484 | *Calypte* | *anna* | 37.7736 | -122.4622 | 1994.00 | 4.25785 | 4.2277 | 0 |
| BLB | 34878 | *Calypte* | *anna* | 37.7167 | -122.5 | 1987.00 | 25.60405 | 0 | 0 |
| BLB | 36231 | *Calypte* | *anna* | 36.5192 | -121.9456 | 1975.00 | 0 | 13.09448 | 0 |
| XC | 42442 | *Calypte* | *anna* | 47.7839 | -122.2739 | 2010.00 | 80.82277 | 19.18344 | 15.17483 |
| XC | 43423 | *Calypte* | *anna* | 47.6976 | -122.2087 | 2010.00 | 39.50789 | 4.67124 | 8.36763 |
| XC | 70079 | *Calypte* | *anna* | 32.2784 | -110.9509 | 2011.00 | 69.19425 | 0 | 0 |
| XC | 80361 | *Calypte* | *anna* | 34.57 | -112.3753 | 2000.00 | 54.42812 | 15.23915 | 15.14803 |
| XC | 132248 | *Calypte* | *anna* | 31.906 | -109.1543 | 2008.00 | 21.12376 | 0 | 0 |
| XC | 132250 | *Calypte* | *anna* | 32.7231 | -116.9473 | 2001.00 | 12.34609 | 0 | 0 |
| XC | 159355 | *Calypte* | *anna* | 48.429528 | -123.479208 | 2012.00 | 6.88827 | 0 | 0 |

| | | | | | | | | | |
|---|---|---|---|---|---|---|---|---|---|
| XC | 169597 | *Calypte* | *anna* | 34.2148 | -118.1477 | 2014.00 | 24.37393 | 0 | 16.03645 |
| XC | 196689 | *Calypte* | *anna* | 31.912 | -109.1426 | 2014.00 | 5.61125 | 0 | 0 |
| XC | 367257 | *Calypte* | *anna* | 39.43 | -122.1862 | 2017.00 | 8.98135 | 0 | 0 |
| XC | 408456 | *Calypte* | *anna* | 34.201 | -118.019 | 2015.00 | 55.99927 | 0 | 2.10983 |
| XC | 442358 | *Calypte* | *anna* | 34.8116 | -115.5608 | 2018.00 | 27.09547 | 0 | 5.58043 |
| XC | 453109 | *Calypte* | *anna* | 32.6287 | -116.923 | 2018.00 | 20.12948 | 13.36248 | 0 |
| XC | 473411 | *Calypte* | *anna* | 47.0562 | -122.9226 | 2019.00 | 8.47885 | 0 | 0 |
| XC | 501895 | *Calypte* | *anna* | 31.403 | -110.2577 | 2019.00 | 44.32586 | 0 | 0 |
| XC | 549602 | *Calypte* | *anna* | 47.7022 | -122.3088 | 2020.00 | 10.33542 | 0 | 0 |
| XC | 624426 | *Calypte* | *anna* | 45.5575 | -122.6912 | 2020.00 | 0 | 0 | 0 |
| BLB | 8 | *Cardinalis* | *cardinalis* | 39.9833 | -82.9167 | 1948.00 | 15.57147 | 0 | 0 |
| BLB | 15 | *Cardinalis* | *cardinalis* | 39.9961 | -83.0189 | 1948.00 | 10.14246 | 0 | 0 |
| BLB | 27 | *Cardinalis* | *cardinalis* | 39.9833 | -82.9167 | 1948.00 | 4.51245 | 0 | 0 |
| BLB | 31 | *Cardinalis* | *cardinalis* | 39.9833 | -82.9167 | 1948.00 | 0 | 6.29532 | 0 |
| BLB | 96 | *Cardinalis* | *cardinalis* | 39.9833 | -82.9167 | 1949.00 | 29.23813 | 4.19085 | 4.80725 |
| BLB | 126 | *Cardinalis* | *cardinalis* | 39.9833 | -82.9167 | 1949.00 | 23.89488 | 0 | 4.85281 |
| BLB | 295 | *Cardinalis* | *cardinalis* | 39.9833 | -82.9167 | 1952.00 | 22.82154 | 4.70273 | 2.21368 |
| BLB | 330 | *Cardinalis* | *cardinalis* | 40.4 | -83.05 | 1953.00 | 27.17319 | 4.59218 | 0 |
| BLB | 331 | *Cardinalis* | *cardinalis* | 40.1 | -83 | 1953.00 | 2.89306 | 5.05247 | 0 |
| BLB | 337 | *Cardinalis* | *cardinalis* | 40.65 | -83 | 1953.00 | 42.50011 | 10.39304 | 0 |
| BLB | 415 | *Cardinalis* | *cardinalis* | 39.9961 | -83.0189 | 1953.00 | 9.6011 | 0 | 10.14715 |
| BLB | 547 | *Cardinalis* | *cardinalis* | 40.0439 | -83.0261 | 1953.00 | 62.15121 | 5.32717 | 0 |
| BLB | 776 | *Cardinalis* | *cardinalis* | 39.9833 | -82.9167 | 1954.00 | 10.32537 | 0 | 0 |
| BLB | 1098 | *Cardinalis* | *cardinalis* | 30.35 | -82.7 | 1954.00 | 5.33856 | 0 | 0 |
| BLB | 1310 | *Cardinalis* | *cardinalis* | 39.35 | -82.25 | 1955.00 | 4.70809 | 0 | 4.16137 |
| BLB | 1352 | *Cardinalis* | *cardinalis* | 39.9403 | -83.0325 | 1955.00 | 15.61234 | 0 | 0 |
| BLB | 1810 | *Cardinalis* | *cardinalis* | 39.6667 | -82.6667 | 1956.00 | 13.59028 | 0 | 0 |
| BLB | 2328 | *Cardinalis* | *cardinalis* | 28.1667 | -81.8333 | 1957.00 | 85.63739 | 9.21116 | 16.0733 |
| BLB | 2435 | *Cardinalis* | *cardinalis* | 39.9403 | -83.0325 | 1957.00 | 6.42865 | 5.69366 | 0 |
| BLB | 3191 | *Cardinalis* | *cardinalis* | 39.9403 | -83.0325 | 1958.00 | 0 | 5.49132 | 5.54157 |
| BLB | 3860 | *Cardinalis* | *cardinalis* | 39.9833 | -82.9167 | 1959.00 | 86.7047 | 5.44308 | 4.69067 |
| BLB | 3886 | *Cardinalis* | *cardinalis* | 39.9403 | -83.0325 | 1959.00 | 3.52487 | 4.39319 | 4.04412 |
| BLB | 4405 | *Cardinalis* | *cardinalis* | 28.0203 | -97.0642 | 1960.00 | 77.19204 | 3.39154 | 6.91574 |
| BLB | 8201 | *Cardinalis* | *cardinalis* | 39.9 | -83.2167 | 1966.00 | 9.74783 | 0 | 0 |
| BLB | 8208 | *Cardinalis* | *cardinalis* | 39.9 | -83.2167 | 1966.00 | 7.21925 | 9.41015 | 0 |
| BLB | 9680 | *Cardinalis* | *cardinalis* | 32.2692 | -110.8758 | 1968.00 | 14.14839 | 0 | 0 |
| BLB | 9700 | *Cardinalis* | *cardinalis* | 32.2692 | -110.8758 | 1968.00 | 7.571 | 0 | 0 |
| BLB | 11276 | *Cardinalis* | *cardinalis* | 40.0833 | -82.9167 | 1971.00 | 10.75484 | 0 | 0 |
| BLB | 11634 | *Cardinalis* | *cardinalis* | 40.0833 | -82.9167 | 1972.00 | 5.31846 | 4.87693 | 5.31377 |

| | | | | | | | | | |
|---|---|---|---|---|---|---|---|---|---|
| BLB | 11876 | *Cardinalis* | *cardinalis* | 35.3333 | -98.1833 | 1972.00 | 5.50874 | 9.75319 | 0 |
| BLB | 13730 | *Cardinalis* | *cardinalis* | 39.9 | -83.2167 | 1976.00 | 5.49668 | 0 | 0 |
| BLB | 14392 | *Cardinalis* | *cardinalis* | 39.9961 | -83.0189 | 1977.00 | 7.18441 | 0 | 0 |
| BLB | 14393 | *Cardinalis* | *cardinalis* | 39.9961 | -83.0189 | 1977.00 | 0 | 3.8927 | 0 |
| BLB | 14394 | *Cardinalis* | *cardinalis* | 39.9961 | -83.0189 | 1977.00 | 8.40314 | 0 | 0 |
| BLB | 15027 | *Cardinalis* | *cardinalis* | 39.9 | -83.2167 | 1979.00 | 11.98563 | 0 | 6.69531 |
| BLB | 15723 | *Cardinalis* | *cardinalis* | 37.9167 | -75.5 | 1981.00 | 8.59275 | 0 | 0 |
| BLB | 15779 | *Cardinalis* | *cardinalis* | 39.9 | -83.2167 | 1982.00 | 12.34877 | 0 | 0 |
| BLB | 16180 | *Cardinalis* | *cardinalis* | 40.0833 | -82.9167 | 1984.00 | 4.6565 | 0 | 0 |
| BLB | 16188 | *Cardinalis* | *cardinalis* | 40.0833 | -82.9167 | 1984.00 | 8.01588 | 0 | 0 |
| BLB | 16287 | *Cardinalis* | *cardinalis* | 40.0833 | -82.9167 | 1985.00 | 9.50864 | 0 | 0 |
| BLB | 16292 | *Cardinalis* | *cardinalis* | 39.9 | -83.2167 | 1985.00 | 14.46597 | 2.27264 | 0 |
| BLB | 16303 | *Cardinalis* | *cardinalis* | 39.9 | -83.2167 | 1985.00 | 14.98991 | 0 | 0 |
| BLB | 16305 | *Cardinalis* | *cardinalis* | 39.9 | -83.2167 | 1985.00 | 7.80282 | 0 | 0 |
| BLB | 21917 | *Cardinalis* | *cardinalis* | 39.9961 | -83.0189 | 1966.00 | 8.17467 | 0 | 0 |
| BLB | 21956 | *Cardinalis* | *cardinalis* | 39.9403 | -83.0325 | 1966.00 | 13.48442 | 0 | 0 |
| BLB | 21960 | *Cardinalis* | *cardinalis* | 39.9403 | -83.0325 | 1966.00 | 6.04072 | 0 | 0 |
| BLB | 21982 | *Cardinalis* | *cardinalis* | 39.9961 | -83.0189 | 1966.00 | 3.96841 | 3.78952 | 0 |
| BLB | 21993 | *Cardinalis* | *cardinalis* | 39.9961 | -83.0189 | 1966.00 | 10.7535 | 0 | 0 |
| BLB | 21997 | *Cardinalis* | *cardinalis* | 39.9961 | -83.0189 | 1966.00 | 5.91476 | 0 | 0 |
| BLB | 22008 | *Cardinalis* | *cardinalis* | 39.9961 | -83.0189 | 1966.00 | 3.82168 | 0 | 0 |
| BLB | 22016 | *Cardinalis* | *cardinalis* | 39.9961 | -83.0189 | 1967.00 | 0 | 0 | 0 |
| BLB | 22026 | *Cardinalis* | *cardinalis* | 39.9961 | -83.0189 | 1967.00 | 0 | 0 | 5.2327 |
| BLB | 22036 | *Cardinalis* | *cardinalis* | 39.9961 | -83.0189 | 1967.00 | 1.5678 | 0 | 9.89523 |
| BLB | 22049 | *Cardinalis* | *cardinalis* | 39.9961 | -83.0189 | 1967.00 | 8.72273 | 0 | 0 |
| BLB | 22055 | *Cardinalis* | *cardinalis* | 39.9961 | -83.0189 | 1967.00 | 11.09654 | 0 | 0 |
| BLB | 22085 | *Cardinalis* | *cardinalis* | 39.9961 | -83.0189 | 1967.00 | 3.83575 | 0 | 4.43272 |
| BLB | 22096 | *Cardinalis* | *cardinalis* | 39.9961 | -83.0189 | 1967.00 | 10.55786 | 0 | 0 |
| BLB | 22118 | *Cardinalis* | *cardinalis* | 39.2961 | -82.6567 | 1967.00 | 0 | 0 | 0 |
| BLB | 22125 | *Cardinalis* | *cardinalis* | 39.9961 | -83.0189 | 1967.00 | 6.77303 | 0 | 0 |
| BLB | 22137 | *Cardinalis* | *cardinalis* | 39.9961 | -83.0189 | 1968.00 | 5.30305 | 0 | 0 |
| BLB | 22581 | *Cardinalis* | *cardinalis* | 53.8333 | -79.1667 | 1957.00 | 11.0148 | 0 | 0 |
| BLB | 32203 | *Cardinalis* | *cardinalis** | 31.8833 | -109.2 | 2001.00 | 17.13994 | 0 | 0 |
| BLB | 32204 | *Cardinalis* | *cardinalis** | 31.8833 | -109.2 | 2001.00 | 19.27523 | 0 | 0 |
| BLB | 5539 | *Cardinalis* | *sinuatus* | 28.1 | -97.4833 | 1962.00 | 18.11479 | 5.00892 | 0 |
| BLB | 5541 | *Cardinalis* | *sinuatus* | 28.1 | -97.4833 | 1962.00 | 59.98376 | 7.27419 | 8.27182 |
| BLB | 5584 | *Cardinalis* | *sinuatus* | 28.1 | -97.4833 | 1962.00 | 47.70802 | 7.54085 | 17.84947 |
| BLB | 9728 | *Cardinalis* | *sinuatus* | 31.8522 | -110.9742 | 1968.00 | 36.75352 | 0 | 0 |
| BLB | 9739 | *Cardinalis* | *sinuatus* | 32.3561 | -111.0878 | 1968.00 | 11.47308 | 0 | 5.86987 |

| | | | | | | | | | |
|---|---|---|---|---|---|---|---|---|---|
| BLB | 10213 | *Cardinalis* | *sinuatus* | 32.3333 | -110.7 | 1969.00 | 25.47742 | 6.61424 | 0 |
| BLB | 10721 | *Cardinalis* | *sinuatus* | 31.6675 | -100.7167 | 1970.00 | 23.92704 | 4.79921 | 0 |
| BLB | 24840 | *Cardinalis* | *sinuatus* | 31.6333 | -111.0333 | 1997.00 | 96.26694 | 8.78906 | 8.7569 |
| BLB | 196 | *Empidonax* | *virescens* | 39.9833 | -82.9167 | 1950.00 | 9.39072 | 4.53523 | 0 |
| BLB | 5160 | *Empidonax* | *virescens* | 40.0833 | -82.9167 | 1961.00 | 16.95636 | 2.12323 | 1.35474 |
| BLB | 6405 | *Empidonax* | *virescens* | 40.0167 | -83.1 | 1963.00 | 4.87291 | 2.42875 | 0 |
| BLB | 10661 | *Empidonax* | *virescens* | 39.5797 | -82.5222 | 1970.00 | 6.38577 | 0 | 0 |
| BLB | 11793 | *Empidonax* | *virescens* | 39.3333 | -84.25 | 1972.00 | 4.62434 | 0 | 2.30413 |
| BLB | 13771 | *Empidonax* | *virescens* | 39.9 | -83.2167 | 1976.00 | 9.1522 | 2.2177 | 2.20966 |
| BLB | 14455 | *Empidonax* | *virescens* | 40.0833 | -82.9167 | 1977.00 | 4.80591 | 0 | 0 |
| BLB | 15125 | *Empidonax* | *virescens* | 40.4 | -83.05 | 1979.00 | 0 | 0 | 2.35103 |
| BLB | 16639 | *Empidonax* | *virescens* | 39.5833 | -83.9833 | 1987.00 | 4.74762 | 5.19451 | 2.37247 |
| XC | 33587 | *Empidonax* | *virescens* | 36.0403 | -93.3406 | 2009.00 | 31.8317 | 0 | 7.61455 |
| XC | 33588 | *Empidonax* | *virescens* | 36.0403 | -93.3406 | 2009.00 | 24.32904 | 2.41066 | 0 |
| XC | 44373 | *Empidonax* | *virescens* | 42.91164591 | -88.47706532 | 2009.00 | 1.72659 | 0 | 0 |
| XC | 182091 | *Empidonax* | *virescens* | 42.6575 | -80.473 | 2001.00 | 2.32624 | 0 | 2.32825 |
| XC | 286544 | *Empidonax* | *virescens* | 29.8235 | -99.5773 | 2015.00 | 18.87524 | 0 | 0 |
| XC | 316116 | *Empidonax* | *virescens* | 34.112 | -91.0917 | 2016.00 | 6.61089 | 0 | 0 |
| XC | 324931 | *Empidonax* | *virescens* | 42.1262 | -79.5069 | 2016.00 | 6.98073 | 4.6498 | 2.37984 |
| XC | 370169 | *Empidonax* | *virescens* | 32.3861 | -85.3675 | 2017.00 | 32.52448 | 3.10478 | 4.15668 |
| XC | 371473 | *Empidonax* | *virescens* | 38.802 | -94.6896 | 2017.00 | 4.75834 | 0 | 1.99459 |
| XC | 371474 | *Empidonax* | *virescens* | 38.802 | -94.6896 | 2017.00 | 9.12875 | 0 | 0 |
| XC | 372010 | *Empidonax* | *virescens* | 38.802 | -94.6896 | 2017.00 | 16.5222 | 2.37046 | 0 |
| XC | 420873 | *Empidonax* | *virescens* | 42.3097 | -76.5008 | 2018.00 | 13.97553 | 0 | 0 |
| XC | 501241 | *Empidonax* | *virescens* | 38.3619 | -77.3422 | 2017.00 | 2.25991 | 0 | 0 |
| BLB | 7184 | *Melozone* | *fusca* | 32.0167 | -109.1667 | 1964.00 | 6.20822 | 3.32387 | 19.66115 |
| BLB | 10118 | *Melozone* | *fusca* | 32.6333 | -110.95 | 1969.00 | 5.99583 | 4.17477 | 25.25163 |
| BLB | 10228 | *Melozone* | *fusca* | 31.5394 | -110.7556 | 1969.00 | 7.16297 | 0 | 21.39243 |
| BLB | 17134 | *Melozone* | *fusca* | 31.85 | -110.9 | 1989.00 | 8.43932 | 1.8626 | 22.66945 |
| BLB | 22880 | *Melozone* | *fusca* | 32.2167 | -110.9167 | ? | 7.36531 | 0 | 14.19998 |
| BLB | 40252 | *Melozone* | *fusca* | 34.1622 | -118.1931 | 1974.00 | 0 | 0 | 3.99856 |
| XC | 18127 | *Melozone* | *fusca* | 32.43619 | -110.90482 | 2008.00 | 3.29305 | 0 | 16.32589 |
| XC | 217984 | *Melozone* | *fusca* | 31.7414 | -110.8873 | 2012.00 | 15.54936 | 0 | 22.23998 |
| XC | 361866 | *Melozone* | *fusca* | 38.2545 | -104.7367 | 2015.00 | 3.34196 | 0 | 10.67511 |
| XC | 530767 | *Melozone* | *fusca* | 19.2553 | -99.0259 | 2020.00 | 0 | 0 | 3.71917 |
| XC | 566741 | *Melozone* | *fusca* | 32.3267 | -110.7033 | 2020.00 | 7.34119 | 0 | 6.94053 |
| XC | 579549 | *Melozone* | *fusca* | 19.1637 | -99.1696 | 2020.00 | 6.58141 | 0 | 20.31641 |
| XC | 1663 | *Myiarchus* | *tuberculifer* | -22.0403 | -64.5584 | 1992 | 8.46143 | 0 | 0 |
| XC | 1721 | *Myiarchus* | *tuberculifer* | -22.1001 | -64.4334 | 1992 | 6.03335 | 0 | 0 |

| | | | | | | | | | |
|---|---|---|---|---|---|---|---|---|---|
| XC | 76805 | *Myiarchus* | *tuberculifer* | 14.5859 | -90.4678 | 2010 | 5.33856 | 0 | 0 |
| XC | 125735 | *Myiarchus* | *tuberculifer* | 14.6724 | -91.485 | 2013 | 8.10365 | 0 | 0 |
| XC | 331718 | *Myiarchus* | *tuberculifer* | 18.0357 | -94.4722 | 2016 | 3.68232 | 0 | 0 |
| XC | 373645 | *Myiarchus* | *tuberculifer* | 13.9475 | -87.1811 | 2017 | 14.49076 | 0 | 2.58687 |
| XC | 429409 | *Myiarchus* | *tuberculifer* | 21.4302 | -87.3345 | 2018 | 9.51936 | 6.73752 | 0 |
| XC | 22439 | *Myiarchus* | *tuberculifer* | 0.1431 | -79.1334 | 2008 | 5.1054 | 0 | 3.9798 |
| XC | 465235 | *Myiarchus* | *tuberculifer* | -3.649 | -79.749 | 2017 | 8.24569 | 0 | 2.52925 |
| XC | 441631 | *Myiarchus* | *tuberculifer* | 28.3714 | -109.0318 | 2018 | 33.13217 | 2.5326 | 0 |
| XC | 352213 | *Myiarchus* | *tuberculifer* | 23.589 | -105.8694 | 2015 | 22.99373 | 7.69897 | 2.73695 |
| XC | 125996 | *Myiarchus* | *tuberculifer* | ? | ? | 2011 | 12.10154 | 0 | 0 |
| BLB | 32001 | *Myiarchus* | *tuberculifer* | 32.2072 | -109.3681 | 1981.00 | 2.34165 | 0 | 2.03077 |
| XC | 11573 | *Myiarchus* | *tuberculifer* | 29.287 | -108.1501 | 2006 | 0 | 0 | 4.19018 |
| XC | 15530 | *Myiarchus* | *tuberculifer* | 10.67 | -85.5 | 1993 | 10.57327 | 0 | 0 |
| XC | 38922 | *Myiarchus* | *tuberculifer* | -5.671292 | -77.758827 | 2006 | 2.05355 | 4.49302 | 0 |
| XC | 41477 | *Myiarchus* | *tuberculifer* | -6.3014 | -79.4623 | 2001 | 6.41592 | 0 | 0 |
| XC | 41478 | *Myiarchus* | *tuberculifer* | -6.3014 | -79.4623 | 2001 | 0 | 0 | 5.94893 |
| XC | 82051 | *Myiarchus* | *tuberculifer* | 4.6834 | -74.3834 | 2009 | 3.66155 | 0 | 0 |
| XC | 219903 | *Myiarchus* | *tuberculifer* | 8.299 | -70.0851 | 2011 | 5.25816 | 1.60532 | 0 |
| XC | 268876 | *Myiarchus* | *tuberculifer* | 13.9641 | -87.1881 | 2015 | 8.00382 | 0 | 2.64181 |
| XC | 279000 | *Myiarchus* | *tuberculifer* | 31.5403 | -110.3754 | 2015 | 1.30248 | 0 | 0 |
| XC | 299054 | *Myiarchus* | *tuberculifer* | -15.8816 | -38.9001 | 2012 | 5.10741 | 2.66727 | 5.03572 |
| XC | 335886 | *Myiarchus* | *tuberculifer* | 12.9006 | -86.5645 | 2014 | 7.16967 | 0 | 0 |
| XC | 360563 | *Myiarchus* | *tuberculifer* | 19.516 | -96.938 | 2016 | 3.10143 | 0 | 0 |
| XC | 382467 | *Myiarchus* | *tuberculifer* | 9.9261 | -75.1074 | 2017 | 2.16075 | 0 | 0 |
| XC | 432855 | *Myiarchus* | *tuberculifer* | -9.5975 | -55.9325 | 2018 | 8.30264 | 0 | 0 |
| XC | 435556 | *Myiarchus* | *tuberculifer* | 6.5574 | -75.8593 | 2017 | 2.9212 | 0 | 0 |
| XC | 504356 | *Myiarchus* | *tuberculifer* | -19.1456 | -40.0645 | 2019 | 9.09927 | 0 | 0 |
| XC | 560657 | *Myiarchus* | *tuberculifer* | 31.9089 | -109.2517 | 2017 | 13.1789 | 0 | 0 |
| XC | 596145 | *Myiarchus* | *tuberculifer* | -3.5761 | -79.5781 | 2019 | 7.83431 | 0 | 0 |
| XC | 672687 | *Myiarchus* | *tuberculifer* | 19.4577 | -103.6175 | 2021.00 | 4.18884 | 0 | 0 |
| XC | 708791 | *Myiarchus* | *tuberculifer* | -9.4642 | -50.0895 | 1999.00 | 5.41561 | 5.42566 | 0 |
| XC | 727101 | *Myiarchus* | *tuberculifer* | 25.766 | -100.2271 | 2022.00 | 6.35093 | 0 | 0 |
| BLB | 7710 | *Passerina* | *amoena* | 44.6125 | -123.2403 | 1965.00 | 24.85432 | 4.20224 | 8.66913 |
| BLB | 9155 | *Passerina* | *amoena* | 47.325 | -114.225 | 1967.00 | 8.56059 | 0 | 0 |
| BLB | 18499 | *Passerina* | *amoena* | 39.6333 | -120.5167 | 1992.00 | 4.03273 | 0 | 3.98985 |
| BLB | 38416 | *Passerina* | *amoena* | 41.7412 | -111.7928 | 1974.00 | 2.47833 | 0 | 3.28032 |
| XC | 1277 | *Passerina* | *amoena* | 38.668892 | -108.304596 | 1991.00 | 3.17044 | 0 | 0 |
| XC | 36565 | *Passerina* | *amoena* | 48.4806 | -121.5834 | 2009.00 | 7.98707 | 0 | 0 |
| XC | 36566 | *Passerina* | *amoena* | 48.4806 | -121.5834 | 2009.00 | 10.12839 | 0 | 0 |

| | | | | | | | | | |
|---|---|---|---|---|---|---|---|---|---|
| XC | 62201 | *Passerina* | *amoena* | 40.137 | -121.448 | 2010.00 | 9.2929 | 0 | 0 |
| XC | 107702 | *Passerina* | *amoena* | 47.014 | -120.706 | 2012.00 | 9.23394 | 0 | 0 |
| XC | 137571 | *Passerina* | *amoena* | 49.0346 | -119.5677 | 2013.00 | 2.45019 | 0 | 0 |
| XC | 137623 | *Passerina* | *amoena* | 49.2997 | -119.5327 | 2013.00 | 3.51147 | 0 | 0 |
| XC | 154997 | *Passerina* | *amoena* | 49.4905 | -119.6205 | 2010.00 | 9.2661 | 0 | 0 |
| XC | 185599 | *Passerina* | *amoena* | 39.9325 | -105.2769 | 2014.00 | 0 | 4.5292 | 0 |
| XC | 251809 | *Passerina* | *amoena* | 49.0804 | -113.8905 | 2015.00 | 6.71541 | 0 | 0 |
| XC | 252848 | *Passerina* | *amoena* | 31.7572 | -110.8438 | 2015.00 | 8.94249 | 0 | 4.08901 |
| XC | 255825 | *Passerina* | *amoena* | 49.0459 | -113.7889 | 2015.00 | 3.94965 | 0 | 0 |
| XC | 269196 | *Passerina* | *amoena* | 46.8437 | -120.7069 | 2015.00 | 4.06355 | 0 | 0 |
| XC | 269198 | *Passerina* | *amoena* | 46.8437 | -120.7069 | 2015.00 | 4.29269 | 0 | 0 |
| XC | 326482 | *Passerina* | *amoena* | 38.7485 | -121.6803 | 2016.00 | 0 | 0 | 3.45184 |
| XC | 331695 | *Passerina* | *amoena* | 40.2286 | -105.2903 | 2016.00 | 3.12555 | 0 | 0 |
| XC | 333586 | *Passerina* | *amoena* | 38.4422 | -119.0056 | 2016.00 | 0 | 4.14328 | 0 |
| XC | 333587 | *Passerina* | *amoena* | 38.4422 | -119.0056 | 2016.00 | 6.39113 | 0 | 0 |
| XC | 376094 | *Passerina* | *amoena* | 46.8897 | -120.7962 | 2017.00 | 3.66825 | 7.23734 | 0 |
| XC | 386101 | *Passerina* | *amoena* | 45.0765 | -108.5295 | 2017.00 | 3.25754 | 0 | 1.90146 |
| XC | 386105 | *Passerina* | *amoena* | 45.0765 | -108.5295 | 2017.00 | 3.4103 | 0 | 0 |
| XC | 415452 | *Passerina* | *amoena* | 44.3896 | -123.3033 | 2018.00 | 2.75705 | 0 | 2.80462 |
| XC | 419780 | *Passerina* | *amoena* | 48.9207 | -122.2617 | 2018.00 | 4.90775 | 0 | 0 |
| XC | 421763 | *Passerina* | *amoena* | 46.6453 | -122.9843 | 2018.00 | 7.42628 | 0 | 0 |
| XC | 468976 | *Passerina* | *amoena* | 38.1378 | -122.6037 | 2019 | 3.0351 | 0 | 0 |
| XC | 478598 | *Passerina* | *amoena* | 48.3113 | -121.5486 | 2019 | 7.78004 | 4.36304 | 0 |
| XC | 478599 | *Passerina* | *amoena* | 48.3114 | -121.5486 | 2019 | 6.92646 | 0 | 0 |
| XC | 483706 | *Passerina* | *amoena* | 38.7985 | -120.3483 | 2019 | 7.59378 | 3.45921 | 0 |
| XC | 548225 | *Passerina* | *amoena* | 33.3543 | -114.6935 | 2020 | 3.44648 | 0 | 0 |
| XC | 559822 | *Passerina* | *amoena* | 32.6675 | -116.9268 | 2020 | 12.62749 | 0 | 0 |
| XC | 563913 | *Passerina* | *amoena* | 40.2286 | -105.2903 | 2020 | 3.81632 | 0 | 0 |
| XC | 582683 | *Passerina* | *amoena* | 38.5431 | -121.6201 | 2020 | 6.68593 | 0 | 0 |
| XC | 642626 | *Passerina* | *amoena* | 48.4991 | -114.1331 | 2017.00 | 6.13385 | 0 | 0 |
| XC | 647816 | *Passerina* | *amoena* | 48.4812 | -123.3732 | 2021.00 | 6.41726 | 0 | 0 |
| XC | 651826 | *Passerina* | *amoena* | 38.7563 | -122.7701 | 2021.00 | 7.63331 | 3.29372 | 0 |
| XC | 662233 | *Passerina* | *amoena* | 44.7361 | -123.1474 | 2021.00 | 6.32614 | 0 | 0 |
| XC | 662824 | *Passerina* | *amoena* | 40.001 | -119.8642 | 2019.00 | 6.85812 | 0 | 0 |
| XC | 665966 | *Passerina* | *amoena* | 40.1864 | -105.3361 | 2021.00 | 6.33686 | 0 | 0 |
| XC | 716430 | *Passerina* | *amoena* | 34.2139 | -118.3069 | 2022.00 | 0 | 0 | 3.53425 |
| XC | 722862 | *Passerina* | *amoena* | 39.3136 | -122.9465 | 2022.00 | 7.38742 | 0 | 0 |
| XC | 56591 | *Poecile* | *carolinensis* | 29.649 | -95.014 | 2010.00 | 23.70259 | 0 | 2.5996 |
| XC | 57321 | *Poecile* | *carolinensis* | 29.649 | -95.014 | 2010.00 | 10.48215 | 3.06123 | 0 |

| | | | | | | | | | |
|---|---|---|---|---|---|---|---|---|---|
| XC | 42453 | *Poecile* | *carolinensis* | 34.1712 | -84.2017 | 2009.00 | 6.76097 | 0 | 0 |
| XC | 734763 | *Poecile* | *carolinensis* | 28.0735 | -82.3744 | 2022.00 | 25.48211 | 3.36675 | 8.09762 |
| BLB | 44185 | *Poecile* | *carolinensis* | 37.6418 | -89.1468 | 2014.00 | 6.50101 | 0 | 0 |
| BLB | 44186 | *Poecile* | *carolinensis* | 35.9786 | -84.9637 | 2014.00 | 3.42906 | 0 | 0 |
| BLB | 175 | *Poecile* | *carolinensis* | 40.1243 | -83.0258 | 1950.00 | 0 | 3.3969 | 0 |
| BLB | 179 | *Poecile* | *carolinensis* | 40.1243 | -83.0258 | 1950.00 | 6.16735 | 3.29975 | 0 |
| BLB | 1312 | *Poecile* | *carolinensis* | 40.0833 | -82.9167 | 1955.00 | 3.35737 | 0 | 2.28537 |
| BLB | 5112 | *Poecile* | *carolinensis* | 39.6667 | -82.6667 | 1961.00 | 3.35402 | 0 | 3.39958 |
| BLB | 11767 | *Poecile* | *carolinensis* | 39.9 | -83.2167 | 1972.00 | 3.40293 | 0 | 0 |
| BLB | 11911 | *Poecile* | *carolinensis* | 39.9 | -83.2167 | 1973.00 | 3.33995 | 2.59491 | 3.32655 |
| BLB | 13003 | *Poecile* | *carolinensis* | 39.5797 | -82.5222 | 1974.00 | 0 | 3.17647 | 0 |
| BLB | 15448 | *Poecile* | *carolinensis* | 39.9 | -83.2167 | 1980.00 | 5.16972 | 0 | 0 |
| BLB | 16251 | *Poecile* | *carolinensis* | 39.2667 | -82.4 | 1984.00 | 9.82622 | 2.28135 | 0 |
| BLB | 16830 | *Poecile* | *carolinensis* | 40.55 | -82.1333 | 1988.00 | 13.25662 | 0 | 0 |
| BLB | 27766 | *Poecile* | *carolinensis* | 39.5797 | -82.5222 | 2002.00 | 2.13663 | 3.0083 | 0 |
| BLB | 40572 | *Poecile* | *carolinensis* | 40.1389 | -83.3128 | 1989.00 | 0 | 3.32387 | 0 |
| XC | 15183 | *Poecile* | *carolinensis* | 35.2176 | -85.922 | 2007.00 | 6.63501 | 0 | 0 |
| XC | 70980 | *Poecile* | *carolinensis* | 29.5826 | -99.7309 | 2010.00 | 8.174 | 0 | 3.06659 |
| XC | 254933 | *Poecile* | *carolinensis* | 32.032 | -97.1212 | 2015.00 | 4.95331 | 2.94666 | 2.95738 |
| XC | 267591 | *Poecile* | *carolinensis* | 32.2517 | -97.8147 | 2015.00 | 8.33547 | 0 | 0 |
| XC | 309925 | *Poecile* | *carolinensis* | 34.3665 | -89.5192 | 2016.00 | 9.26275 | 0 | 3.30377 |
| XC | 310196 | *Poecile* | *carolinensis* | 38.0915 | -78.4939 | 2016.00 | 0 | 0 | 3.36675 |
| XC | 385887 | *Poecile* | *carolinensis* | 39.9826 | -74.2374 | 2017 | 1.26831 | 0 | 0 |
| XC | 416023 | *Poecile* | *carolinensis* | 38.7877 | -77.2332 | 2018.00 | 6.40922 | 0 | 3.17781 |
| XC | 417997 | *Poecile* | *carolinensis* | 40.5652 | -83.6255 | 2018.00 | 9.96692 | 0 | 0 |
| XC | 434624 | *Poecile* | *carolinensis* | 34.9961 | -82.4037 | 2018.00 | 14.85658 | 0 | 0 |
| XC | 458045 | *Poecile* | *carolinensis* | 37.2602 | -97.4172 | 2019.00 | 12.8506 | 0 | 0 |
| XC | 483022 | *Poecile* | *carolinensis* | 34.0091 | -86.1929 | 2019 | 3.54296 | 0 | 1.85992 |
| XC | 499562 | *Poecile* | *carolinensis* | 38.3619 | -77.3422 | 2017 | 11.94744 | 0 | 3.01232 |
| XC | 616885 | *Poecile* | *carolinensis* | 37.9754 | -90.8983 | 2021.00 | 5.71443 | 0 | 0 |
| XC | 438079 | *Vireo* | *altiloquus* | 18.4485 | -69.0962 | 2018.00 | 10.06072 | 2.5661 | 5.58713 |
| XC | 489358 | *Vireo* | *altiloquus* | 25.2792 | -80.2983 | 2019.00 | 16.482 | 5.20255 | 0 |
| XC | 495838 | *Vireo* | *altiloquus* | 13.211 | -59.5969 | 2019.00 | 35.63596 | 0 | 1.5812 |
| XC | 82985 | *Vireo* | *altiloquus* | 22.6689 | -83.4789 | 2011.00 | 2.54332 | 3.74932 | 0 |
| XC | 82986 | *Vireo* | *altiloquus* | 22.6689 | -83.4789 | 2011.00 | 23.52906 | 0 | 5.53889 |
| XC | 82987 | *Vireo* | *altiloquus* | 21.01 | -77.7198 | 2011.00 | 15.47298 | 2.50178 | 2.54332 |
| XC | 104633 | *Vireo* | *altiloquus* | 24.6959 | -81.3209 | 2012.00 | 39.58025 | 7.33047 | 5.04845 |
| XC | 104634 | *Vireo* | *altiloquus* | 24.6959 | -81.3209 | 2012.00 | 39.8114 | 2.58754 | 5.4337 |
| XC | 256744 | *Vireo* | *altiloquus* | 22.657 | -83.445 | 2015.00 | 23.99806 | 2.86961 | 2.90378 |

| | | | | | | | | | |
|---|---|---|---|---|---|---|---|---|---|
| XC | 97198 | *Vireo* | *altiloquus* | 18.492755 | -69.952826 | 2012.00 | 16.25353 | 2.61769 | 2.55002 |
| XC | 215245 | *Vireo* | *altiloquus* | 17.6285 | -63.2493 | 2013.00 | 4.73422 | 0 | 1.67969 |
| XC | 370027 | *Vireo* | *altiloquus* | 18.0442 | -66.114 | 2017.00 | 2.85152 | 0 | 0 |
| BLB | 13760 | *Zonotrichia* | *leucophrys* | 40.0833 | -82.9167 | 1976.00 | 11.39603 | 3.72587 | 16.78618 |
| BLB | 18611 | *Zonotrichia* | *leucophrys* | 61.75 | -150.0833 | 1993.00 | 55.73261 | 7.37134 | 7.05979 |
| BLB | 18613 | *Zonotrichia* | *leucophrys* | 61.5294 | -149.9075 | 1993.00 | 29.48201 | 12.71593 | 3.72453 |
| BLB | 18799 | *Zonotrichia* | *leucophrys* | 61.25 | -149.25 | 1993.00 | 43.81264 | 3.75937 | 14.9678 |
| BLB | 23095 | *Zonotrichia* | *leucophrys* | 51.1667 | -80.3333 | 1958.00 | 17.50576 | 4.2076 | 0 |
| BLB | 23096 | *Zonotrichia* | *leucophrys* | 51.1667 | -80.3333 | 1958.00 | 23.2691 | 7.7385 | 0 |
| BLB | 23097 | *Zonotrichia* | *leucophrys* | 58.725 | -94.1167 | 1959.00 | 20.67285 | 0 | 0 |
| BLB | 23875 | *Zonotrichia* | *leucophrys* | 38.5 | -121.75 | 1997.00 | 34.72744 | 4.16539 | 12.55111 |
| BLB | 23876 | *Zonotrichia* | *leucophrys* | 38.5 | -121.75 | 1997.00 | 39.39265 | 4.38984 | 4.08968 |
| BLB | 23877 | *Zonotrichia* | *leucophrys* | 38.5 | -121.75 | 1997.00 | 20.84839 | 0 | 4.20358 |
| BLB | 23903 | *Zonotrichia* | *leucophrys* | 58.7333 | -93.8167 | 1997.00 | 12.08814 | 4.11112 | 4.11112 |
| BLB | 24704 | *Zonotrichia* | *leucophrys* | 38.0833 | -122.9167 | 1998.00 | 80.84354 | 12.65965 | 39.46367 |
| BLB | 29924 | *Zonotrichia* | *leucophrys* | 38.5 | -121.75 | 1990.00 | 12.95646 | 4.221 | 0 |
| BLB | 33624 | *Zonotrichia* | *leucophrys* | 33.2167 | -116.25 | 1976.00 | 30.53056 | 0 | 4.01196 |
| BLB | 35099 | *Zonotrichia* | *leucophrys* | 60.5711 | -151.2478 | 1968.00 | 7.82359 | 0 | 10.29321 |
| BLB | 35101 | *Zonotrichia* | *leucophrys* | 61.5833 | -149.1 | 1968.00 | 11.27878 | 3.50946 | 0 |
| BLB | 35102 | *Zonotrichia* | *leucophrys* | 61.0561 | -149.7972 | 1968.00 | 11.10056 | 3.46524 | 0 |
| BLB | 35130 | *Zonotrichia* | *leucophrys* | 34.4125 | -119.8481 | 1969.00 | 6.36835 | 0 | 0 |
| BLB | 35608 | *Zonotrichia* | *leucophrys* | 37.8775 | -122.2508 | 1969.00 | 6.27254 | 0 | 7.102 |
| BLB | 37698 | *Zonotrichia* | *leucophrys* | 49.0878 | -120.8166 | 1970.00 | 21.85808 | 0 | 0 |
| BLB | 37953 | *Zonotrichia* | *leucophrys* | 37.2502 | -119.7513 | 1987.00 | 14.6998 | 0 | 5.49266 |
| BLB | 41400 | *Zonotrichia* | *leucophrys* | 63.2941 | -148.5528 | 1996.00 | 12.18797 | 0 | 8.19678 |
| BLB | 42491 | *Zonotrichia* | *leucophrys* | 34.4423 | -119.8112 | 1967.00 | 6.28527 | 0 | 0 |
| BLB | 42554 | *Zonotrichia* | *leucophrys* | 64.8548 | -147.8379 | 1973.00 | 8.44736 | 0 | 8.42994 |
| BLB | 2529 | *Zonotrichia* | *leucophrys* | 39.6667 | -82.6667 | 1957.00 | 57.72117 | 0 | 4.44746 |
| BLB | 2570 | *Zonotrichia* | *leucophrys* | 39.6667 | -82.6667 | 1957.00 | 51.33607 | 3.90476 | 3.57311 |
| BLB | 5132 | *Zonotrichia* | *leucophrys* | 39.9403 | -83.0325 | 1961.00 | 63.27949 | 0 | 4.14998 |
| BLB | 5769 | *Zonotrichia* | *leucophrys* | 39.9403 | -83.0325 | 1962.00 | 15.97414 | 4.15735 | 4.24847 |
| BLB | 7412 | *Zonotrichia* | *leucophrys* | 39.9833 | -82.9167 | 1965.00 | 70.65351 | 10.21415 | 33.76398 |
| BLB | 8296 | *Zonotrichia* | *leucophrys* | 41.5333 | -82.9333 | 1966.00 | 27.97853 | 0 | 0 |
| BLB | 8774 | *Zonotrichia* | *leucophrys* | 39.6667 | -82.6667 | 1967.00 | 39.12465 | 8.50096 | 7.60048 |
| BLB | 9398 | *Zonotrichia* | *leucophrys* | 40.0833 | -82.9167 | 1968.00 | 33.71038 | 0 | 4.10643 |
| BLB | 23900 | *Zonotrichia* | *leucophrys* | 58.7333 | -93.8167 | 1997.00 | 15.90111 | 11.96955 | 3.7654 |
| BLB | 25372 | *Zonotrichia* | *leucophrys* | 40 | -83 | 1999.00 | 0 | 0 | 7.76932 |
| BLB | 26743 | *Zonotrichia* | *leucophrys* | 40 | -83 | 2001.00 | 9.91466 | 0 | 0 |
| BLB | 21132 | *Zonotrichia* | *leucophrys* | 39.6333 | -123.7833 | 2003.00 | 35.5301 | 3.77545 | 15.35037 |

| | | | | | | | | | |
|---|---|---|---|---|---|---|---|---|---|
| BLB | 21133 | *Zonotrichia* | *leucophrys* | 39.6 | -123.7833 | 2003.00 | 0 | 4.0602 | 4.11849 |
| BLB | 22428 | *Zonotrichia* | *leucophrys* | 39.3 | -123.7833 | 2003.00 | 8.23631 | 0 | 0 |
| BLB | 22887 | *Zonotrichia* | *leucophrys* | 37.7289 | -122.4948 | 1971.00 | 11.79334 | 0 | 3.94697 |
| BLB | 23083 | *Zonotrichia* | *leucophrys* | 39.0333 | -123.7 | 2003.00 | 3.67696 | 3.03309 | 3.15436 |
| BLB | 23326 | *Zonotrichia* | *leucophrys* | 39 | -123.6833 | 2003.00 | 15.26461 | 3.97645 | 3.58919 |
| BLB | 23537 | *Zonotrichia* | *leucophrys* | 39 | -123.6833 | 2003.00 | 8.47148 | 0 | 0 |
| BLB | 23882 | *Zonotrichia* | *leucophrys* | 37.7289 | -122.4948 | 1971.00 | 3.98315 | 7.68155 | 0 |
| BLB | 23941 | *Zonotrichia* | *leucophrys* | 36.75 | -121.75 | 1997.00 | 3.87059 | 0 | 0 |
| BLB | 23948 | *Zonotrichia* | *leucophrys* | 36.7833 | -121.75 | 1997.00 | 35.53948 | 0 | 0 |
| BLB | 24518 | *Zonotrichia* | *leucophrys* | 37.7289 | -122.4948 | 1971.00 | 11.17627 | 0 | 0 |
| BLB | 24688 | *Zonotrichia* | *leucophrys* | 38.0833 | -122.9167 | 1998.00 | 10.63089 | 6.15931 | 0 |
| BLB | 24705 | *Zonotrichia* | *leucophrys* | 38.0833 | -122.9167 | 1998.00 | 20.23735 | 3.8592 | 0 |
| BLB | 24706 | *Zonotrichia* | *leucophrys* | 38.0833 | -122.9167 | 1998.00 | 10.18668 | 0 | 0 |
| BLB | 24707 | *Zonotrichia* | *leucophrys* | 38.0833 | -122.9167 | 1998.00 | 7.59914 | 3.83441 | 7.24538 |
| BLB | 24709 | *Zonotrichia* | *leucophrys* | 38.0833 | -122.9167 | 1998.00 | 4.41731 | 0 | 0 |
| BLB | 24710 | *Zonotrichia* | *leucophrys* | 38.0833 | -122.9167 | 1998.00 | 36.64565 | 0 | 7.93548 |
| BLB | 24711 | *Zonotrichia* | *leucophrys* | 38.0833 | -122.9167 | 1998.00 | 3.38953 | 0 | 0 |
| BLB | 24713 | *Zonotrichia* | *leucophrys* | 38.0833 | -122.9167 | 1998.00 | 19.58477 | 5.48328 | 0 |
| BLB | 24714 | *Zonotrichia* | *leucophrys* | 38.0833 | -122.9167 | 1998.00 | 3.50477 | 0 | 0 |
| BLB | 26976 | *Zonotrichia* | *leucophrys* | 37.8569 | -122.2978 | 1971.00 | 4.20291 | 4.04144 | 0 |
| BLB | 28219 | *Zonotrichia* | *leucophrys* | 38.5667 | -123.3333 | 2003.00 | 4.21631 | 0 | 8.52508 |
| BLB | 29612 | *Zonotrichia* | *leucophrys* | 38.3022 | -123.0572 | 1993.00 | 8.27651 | 3.99454 | 0 |
| BLB | 32507 | *Zonotrichia* | *leucophrys* | 37.6139 | -122.4869 | 1981.00 | 7.43633 | 0 | 0 |
| BLB | 33184 | *Zonotrichia* | *leucophrys* | 37.7667 | -122.45 | 1981.00 | 24.24462 | 7.08927 | 9.45102 |
| BLB | 33185 | *Zonotrichia* | *leucophrys* | 37.7667 | -122.45 | 1981.00 | 56.16275 | 3.1088 | 0 |
| BLB | 33261 | *Zonotrichia* | *leucophrys* | 37.9667 | -119.1 | 1982.00 | 4.17477 | 0 | 0 |
| BLB | 33332 | *Zonotrichia* | *leucophrys* | 37.7289 | -122.4948 | 1974.00 | 16.01032 | 3.55368 | 10.42118 |
| BLB | 33407 | *Zonotrichia* | *leucophrys* | 37.7667 | -122.4667 | 1981.00 | 3.96238 | 0 | 0 |
| BLB | 33463 | *Zonotrichia* | *leucophrys* | 37.7125 | -122.4992 | 1990.00 | 21.40717 | 4.00124 | 4.00124 |
| BLB | 33665 | *Zonotrichia* | *leucophrys* | 37.7289 | -122.4948 | 1969.00 | 20.71774 | 0 | 0 |
| BLB | 33861 | *Zonotrichia* | *leucophrys* | 38.0667 | -122.8844 | 1985.00 | 7.68155 | 0 | 13.7819 |
| BLB | 33944 | *Zonotrichia* | *leucophrys* | 37.7667 | -122.45 | 1982.00 | 10.69588 | 3.95501 | 3.95501 |
| BLB | 34043 | *Zonotrichia* | *leucophrys* | 37.7669 | -122.4894 | 1991.00 | 3.15704 | 3.1624 | 0 |
| BLB | 34281 | *Zonotrichia* | *leucophrys* | 37.803 | -122.4745 | 1992.00 | 19.25714 | 0 | 7.81488 |
| BLB | 34417 | *Zonotrichia* | *leucophrys* | 37.7667 | -122.4667 | 1993.00 | 17.84612 | 0 | 0 |
| BLB | 34570 | *Zonotrichia* | *leucophrys* | 37.7667 | -122.4667 | 1992.00 | 15.18019 | 0 | 10.95383 |
| BLB | 34823 | *Zonotrichia* | *leucophrys* | 36.5217 | -121.9528 | 1976.00 | 17.6813 | 0 | 0 |
| BLB | 35222 | *Zonotrichia* | *leucophrys* | 37.7667 | -122.4667 | 1968.00 | 13.06165 | 0 | 6.64908 |
| BLB | 35290 | *Zonotrichia* | *leucophrys* | 37.7667 | -122.4667 | 1969.00 | 9.75855 | 0 | 0 |

| | | | | | | | | | |
|---|---|---|---|---|---|---|---|---|---|
| BLB | 35490 | *Zonotrichia* | *leucophrys* | 32.5839 | -117.1131 | 1968.00 | 3.93625 | 0 | 0 |
| BLB | 35513 | *Zonotrichia* | *leucophrys* | 37.93 | -122.7353 | 1969.00 | 6.83132 | 0 | 0 |
| BLB | 35517 | *Zonotrichia* | *leucophrys* | 37.3192 | -122.2742 | 1969.00 | 0 | 0 | 6.22564 |
| BLB | 35525 | *Zonotrichia* | *leucophrys* | 37.9058 | -122.2386 | 1969.00 | 6.36701 | 0 | 0 |
| BLB | 35732 | *Zonotrichia* | *leucophrys* | 37.8 | -122.4636 | 1969.00 | 0 | 2.99557 | 3.09674 |
| BLB | 35795 | *Zonotrichia* | *leucophrys* | 37.9158 | -122.3117 | 1970.00 | 10.16658 | 0 | 0 |
| BLB | 35806 | *Zonotrichia* | *leucophrys* | 38.041 | -122.7994 | 1970.00 | 0 | 0 | 0 |
| BLB | 35896 | *Zonotrichia* | *leucophrys* | 37.7289 | -122.4948 | 1970.00 | 17.96471 | 0 | 0 |
| BLB | 35955 | *Zonotrichia* | *leucophrys* | 37.7667 | -122.45 | 1971.00 | 2.57816 | 0 | 0 |
| BLB | 36059 | *Zonotrichia* | *leucophrys* | 37.7667 | -122.4667 | 1971.00 | 12.84725 | 4.48498 | 4.48498 |
| BLB | 36060 | *Zonotrichia* | *leucophrys* | 37.7667 | -122.4667 | 1971.00 | 15.72423 | 0 | 15.84416 |
| BLB | 36166 | *Zonotrichia* | *leucophrys* | 37.8569 | -122.2978 | 1971.00 | 12.04459 | 0 | 0 |
| BLB | 37006 | *Zonotrichia* | *leucophrys* | 35.068 | -120.6093 | 1982.00 | 18.72114 | 0 | 0 |
| BLB | 38164 | *Zonotrichia* | *leucophrys* | 37.874 | -122.239 | 1970.00 | 3.90409 | 3.8793 | 7.38072 |
| BLB | 39486 | *Zonotrichia* | *leucophrys* | 37.7667 | -122.4667 | 1995.00 | 42.66962 | 10.96254 | 3.74999 |
| BLB | 39997 | *Zonotrichia* | *leucophrys* | 38.3174 | -123.0711 | 1995.00 | 49.03261 | 0 | 0 |
| BLB | 41913 | *Zonotrichia* | *leucophrys* | 37.7667 | -122.4667 | 1994.00 | 7.27687 | 4.13993 | 3.67294 |
| BLB | 42360 | *Zonotrichia* | *leucophrys* | 37.7883 | -122.4608 | 1977.00 | 8.24301 | 0 | 8.02392 |
| BLB | 42500 | *Zonotrichia* | *leucophrys* | 37.7667 | -122.4667 | 1992.00 | 47.73147 | 3.7453 | 3.68232 |
| XC | 445848 | *Zonotrichia* | *leucophrys* | 39.3081 | -123.81 | 2015.00 | 11.77592 | 3.69974 | 0 |
| XC | 445849 | *Zonotrichia* | *leucophrys* | 39.2263 | -123.7203 | 2015.00 | 24.39135 | 4.13658 | 8.22894 |
| XC | 459465 | *Zonotrichia* | *leucophrys* | 37.771 | -122.4924 | 2019.00 | 11.66336 | 3.79354 | 0 |
| XC | 610641 | *Zonotrichia* | *leucophrys* | 37.9085 | -122.3511 | 2020.00 | 10.1773 | 0 | 3.57579 |
| BLB | 6596 | *Zonotrichia* | *leucophrys* | 41.3333 | -106.3 | 1963.00 | 11.03892 | 0 | 3.89203 |
| BLB | 6597 | *Zonotrichia* | *leucophrys* | 43.0167 | -110.1167 | 1963.00 | 36.11233 | 4.05283 | 0 |
| BLB | 20614 | *Zonotrichia* | *leucophrys* | 38.3333 | -119.6333 | 1993.00 | 6.38711 | 0 | 0 |
| BLB | 20616 | *Zonotrichia* | *leucophrys* | 38.3333 | -119.6333 | 1993.00 | 37.4597 | 10.85802 | 29.12691 |
| BLB | 20617 | *Zonotrichia* | *leucophrys* | 38.3333 | -119.6333 | 1993.00 | 92.85597 | 4.07092 | 4.27929 |
| BLB | 20628 | *Zonotrichia* | *leucophrys* | 38.7 | -119.9833 | 1993.00 | 118.01246 | 14.81236 | 15.68336 |
| BLB | 32036 | *Zonotrichia* | *leucophrys* | 37.9 | -119.1167 | 1980.00 | 10.81581 | 0 | 0 |
| BLB | 33278 | *Zonotrichia* | *leucophrys* | 37.9 | -119.1167 | 1984.00 | 23.70192 | 0 | 0 |
| BLB | 33894 | *Zonotrichia* | *leucophrys* | 37.9778 | -119.1019 | 1988.00 | 3.84513 | 6.00387 | 10.84462 |
| BLB | 33967 | *Zonotrichia* | *leucophrys* | 42.4167 | -119.6667 | 1975.00 | 7.76195 | 0 | 0 |
| BLB | 35008 | *Zonotrichia* | *leucophrys* | 37.9667 | -119.1 | 1989.00 | 7.46045 | 0 | 10.41448 |
| BLB | 7718 | *Zonotrichia* | *leucophrys* | 44.5833 | -123.3167 | 1965.00 | 55.007 | 7.64403 | 7.74118 |
| BLB | 24672 | *Zonotrichia* | *leucophrys* | 40 | -83 | 1998.00 | 3.86925 | 0 | 4.05216 |
| BLB | 24739 | *Zonotrichia* | *leucophrys* | 46.25 | -124.0833 | 1998.00 | 99.75094 | 8.13849 | 29.3192 |
| BLB | 24773 | *Zonotrichia* | *leucophrys* | 46.25 | -124.0833 | 1998.00 | 2.82405 | 0 | 0 |
| BLB | 24869 | *Zonotrichia* | *leucophrys* | 46.2167 | -124 | 1998.00 | 11.6848 | 0 | 0 |

| | | | | | | | | | |
|---|---|---|---|---|---|---|---|---|---|
| BLB | 24878 | *Zonotrichia* | *leucophrys* | 46.25 | -124.0833 | 1998.00 | 11.62919 | 3.61465 | 0 |
| BLB | 24880 | *Zonotrichia* | *leucophrys* | 46.25 | -124.0833 | 1998.00 | 19.42531 | 3.90476 | 0 |
| BLB | 24881 | *Zonotrichia* | *leucophrys* | 46.2286 | -124.01 | 1998.00 | 3.44715 | 3.64212 | 0 |
| BLB | 24882 | *Zonotrichia* | *leucophrys* | 46.2167 | -124 | 1998.00 | 60.43936 | 8.19477 | 32.07826 |
| BLB | 24883 | *Zonotrichia* | *leucophrys* | 46.2167 | -124 | 1998.00 | 123.72957 | 20.11541 | 22.4785 |
| BLB | 24894 | *Zonotrichia* | *leucophrys* | 46.2286 | -124.01 | 1998.00 | 76.87044 | 12.1672 | 41.55943 |
| BLB | 25949 | *Zonotrichia* | *leucophrys* | 48.5 | -124.5 | 1999.00 | 50.6989 | 3.98516 | 11.76989 |
| BLB | 26285 | *Zonotrichia* | *leucophrys* | 48.2 | -122.7 | 2000.00 | 13.8623 | 4.42468 | 0 |
| BLB | 26355 | *Zonotrichia* | *leucophrys* | 46.2847 | -124.0769 | 2000.00 | 7.80483 | 3.71984 | 0 |
| BLB | 26845 | *Zonotrichia* | *leucophrys* | 48 | -123.75 | 2001.00 | 12.13303 | 4.27259 | 0 |
| BLB | 29176 | *Zonotrichia* | *leucophrys* | 40.4833 | -124.1 | 2004.00 | 10.99068 | 0 | 0 |
| BLB | 29177 | *Zonotrichia* | *leucophrys* | 41.75 | -124.2 | 2004.00 | 4.00325 | 0 | 3.5309 |
| BLB | 31650 | *Zonotrichia* | *leucophrys* | 41.9283 | -124.1458 | 2005.00 | 0 | 3.7654 | 3.60862 |
| BLB | 33235 | *Zonotrichia* | *leucophrys* | 37.7667 | -122.45 | 1980.00 | 23.87277 | 2.04082 | 0 |
| BLB | 33834 | *Zonotrichia* | *leucophrys* | 47.0072 | -122.9094 | 1977.00 | 8.36562 | 0 | 3.90744 |
| BLB | 35011 | *Zonotrichia* | *leucophrys* | 43.9667 | -124.0833 | 1985.00 | 46.79682 | 0 | 13.45896 |
| BLB | 35043 | *Zonotrichia* | *leucophrys* | 34.4 | -119.8333 | 1968.00 | 10.8875 | 0 | 7.96362 |
| BLB | 35047 | *Zonotrichia* | *leucophrys* | 34.4 | -119.8333 | 1968.00 | 3.9061 | 3.22404 | 0 |
| BLB | 35110 | *Zonotrichia* | *leucophrys* | 34.4631 | -119.8319 | 1968.00 | 0 | 3.92285 | 3.94831 |
| BLB | 35123 | *Zonotrichia* | *leucophrys* | 34.4 | -119.8333 | 1968.00 | 11.61847 | 3.95099 | 0 |
| BLB | 35124 | *Zonotrichia* | *leucophrys* | 34.4 | -119.8333 | 1969.00 | 10.47679 | 0 | 0 |
| BLB | 35144 | *Zonotrichia* | *leucophrys* | 34.4 | -119.8333 | 1970.00 | 6.82663 | 3.47931 | 3.47931 |
| BLB | 36565 | *Zonotrichia* | *leucophrys* | 40.7891 | -124.176 | 1970.00 | 24.74846 | 0 | 0 |
| BLB | 36712 | *Zonotrichia* | *leucophrys* | 43.0667 | -124.4333 | 1970.00 | 13.80133 | 0 | 0 |
| BLB | 36819 | *Zonotrichia* | *leucophrys* | 44.6378 | -124.0339 | 1970.00 | 11.58564 | 0 | 3.57244 |
| BLB | 36820 | *Zonotrichia* | *leucophrys* | 45.4627 | -123.8366 | 1970.00 | 17.43273 | 3.63341 | 0 |
| BLB | 36893 | *Zonotrichia* | *leucophrys* | 46.25 | -124.0833 | 1970.00 | 0 | 0 | 3.83374 |
| BLB | 36961 | *Zonotrichia* | *leucophrys* | 34.4124 | -119.8518 | 1982.00 | 29.71182 | 3.6582 | 10.18199 |
| BLB | 37080 | *Zonotrichia* | *leucophrys* | 47.5507 | -122.0442 | 1970.00 | 11.10458 | 3.82168 | 7.68356 |
| BLB | 37085 | *Zonotrichia* | *leucophrys* | 47.3045 | -122.5201 | 1970.00 | 11.74644 | 0 | 3.73458 |
| BLB | 37704 | *Zonotrichia* | *leucophrys* | 46.707 | -123.98 | 1970.00 | 16.5088 | 0 | 16.57848 |
| BLB | 38159 | *Zonotrichia* | *leucophrys* | 42.8411 | -124.5419 | 1970.00 | 11.86034 | 0 | 7.64872 |
| BLB | 38431 | *Zonotrichia* | *leucophrys* | 42.7329 | -119.5507 | 1977.00 | 8.04 | 0 | 0 |
| BLB | 39290 | *Zonotrichia* | *leucophrys* | 34.4243 | -119.7511 | 1986.00 | 4.76772 | 0 | 0 |
| BLB | 39650 | *Zonotrichia* | *leucophrys* | 36.9664 | -95.844 | 1983.00 | 2.9346 | 0 | 0 |
| BLB | 40686 | *Zonotrichia* | *leucophrys* | 48.7376 | -122.7929 | 1977.00 | 31.35466 | 0 | 16.44314 |
| BLB | 42570 | *Zonotrichia* | *leucophrys* | 34.4631 | -119.8319 | 1967.00 | 3.56172 | 0 | 6.499 |
| XC | 160205 | *Zonotrichia* | *leucophrys* | 48.395338 | -123.599262 | 2013.00 | 7.70165 | 0 | 0 |
| XC | 182564 | *Zonotrichia* | *leucophrys* | 48.5093 | -123.4537 | 2014.00 | 0 | 0 | 4.17075 |

| | | | | | | | | | |
|---|---|---|---|---|---|---|---|---|---|
| XC | 253833 | *Zonotrichia* | *leucophrys* | 44.3799 | -121.6917 | 2015.00 | 7.50199 | 0 | 3.14096 |
| XC | 253837 | *Zonotrichia* | *leucophrys* | 43.661 | -124.207 | 2015.00 | 7.9864 | 3.39623 | 0 |
| XC | 253839 | *Zonotrichia* | *leucophrys* | 44.753 | -123.3028 | 2015.00 | 7.035 | 0 | 0 |
| XC | 570694 | *Zonotrichia* | *leucophrys* | 48.4269 | -123.3797 | 2020.00 | 3.79354 | 3.69237 | 0 |
| BLB | 218 | *Zonotrichia* | *leucophrys* | 39.9961 | -83.0189 | 1951.00 | 18.97038 | 0 | 0 |
| BLB | 1378 | *Zonotrichia* | *leucophrys* | 39.6667 | -82.6667 | 1955.00 | 41.55809 | 8.49024 | 0 |
| BLB | 11529 | *Zonotrichia* | *leucophrys* | 39.3264 | -119.8778 | 1971.00 | 11.8389 | 0 | 8.13581 |
| BLB | 11538 | *Zonotrichia* | *leucophrys* | 36.6167 | -121.8167 | 1971.00 | 23.55251 | 3.81431 | 4.13457 |
| BLB | 11543 | *Zonotrichia* | *leucophrys* | 36.5 | -121.8333 | 1971.00 | 31.85515 | 5.71778 | 0 |
| BLB | 11545 | *Zonotrichia* | *leucophrys* | 36.5 | -121.8333 | 1971.00 | 3.75334 | 0 | 8.14586 |
| BLB | 12866 | *Zonotrichia* | *leucophrys* | 40.0833 | -82.9167 | 1974.00 | 8.33614 | 0 | 0 |
| BLB | 14778 | *Zonotrichia* | *leucophrys* | 41.5333 | -82.9333 | 1978.00 | 15.50246 | 0 | 4.10978 |
| BLB | 18009 | *Zonotrichia* | *leucophrys* | 58.725 | -94.1167 | 1989.00 | 15.66192 | 0 | 0 |
| BLB | 18605 | *Zonotrichia* | *leucophrys* | 61.5294 | -149.9075 | 1993.00 | 62.91501 | 4.21564 | 10.66439 |
| BLB | 18630 | *Zonotrichia* | *leucophrys* | 63.3333 | -150.5006 | 1993.00 | 4.1674 | 4.12184 | 0 |
| BLB | 18749 | *Zonotrichia* | *leucophrys* | 39.5833 | -120.4833 | 1992.00 | 20.00486 | 0 | 0 |
| BLB | 20613 | *Zonotrichia* | *leucophrys* | 38.3333 | -119.6333 | 1993.00 | 3.90811 | 3.93558 | 0 |
| BLB | 21131 | *Zonotrichia* | *leucophrys* | 39.6333 | -123.7833 | 2003.00 | 13.28141 | 0 | 0 |
| BLB | 23093 | *Zonotrichia* | *leucophrys* | 58.725 | -94.1167 | 1959.00 | 16.51885 | 4.17075 | 4.16539 |
| BLB | 23094 | *Zonotrichia* | *leucophrys* | 51.1667 | -80.3333 | 1958.00 | 29.55705 | 0 | 0 |
| BLB | 23681 | *Zonotrichia* | *leucophrys* | 37.9 | -119.1167 | 1990.00 | 27.52025 | 3.63475 | 0 |
| BLB | 23863 | *Zonotrichia* | *leucophrys* | 58.7333 | -93.8167 | 1997.00 | 19.83401 | 0 | 0 |
| BLB | 23874 | *Zonotrichia* | *leucophrys* | 38.5 | -121.75 | 1997.00 | 50.90928 | 9.21317 | 10.93507 |
| BLB | 23901 | *Zonotrichia* | *leucophrys* | 58.7333 | -93.8167 | 1997.00 | 19.91374 | 4.24981 | 0 |
| BLB | 23902 | *Zonotrichia* | *leucophrys* | 58.7333 | -93.8167 | 1997.00 | 27.02847 | 0 | 4.10643 |
| BLB | 23928 | *Zonotrichia* | *leucophrys* | 38.5 | -121.75 | 1991.00 | 49.57933 | 3.53023 | 2.34433 |
| BLB | 23939 | *Zonotrichia* | *leucophrys* | 36.75 | -121.75 | 1997.00 | 6.40654 | 0 | 3.417 |
| BLB | 24673 | *Zonotrichia* | *leucophrys* | 38.5 | -121.75 | 1998.00 | 39.74976 | 3.66624 | 0 |
| BLB | 24692 | *Zonotrichia* | *leucophrys* | 40 | -83 | 1998.00 | 11.46571 | 0 | 0 |
| BLB | 24738 | *Zonotrichia* | *leucophrys* | 46.2286 | -124.01 | 1998.00 | 14.82174 | 4.69402 | 2.16812 |
| BLB | 27439 | *Zonotrichia* | *leucophrys* | 38.7 | -119.9833 | 1995.00 | 12.91157 | 0 | 3.6582 |
| BLB | 30928 | *Zonotrichia* | *leucophrys* | 48.15 | -123.2 | 1996.00 | 10.10561 | 0 | 0 |
| BLB | 34996 | *Zonotrichia* | *leucophrys* | 39.6833 | -120.4 | 1989.00 | 10.69454 | 0 | 3.47931 |
| BLB | 35122 | *Zonotrichia* | *leucophrys* | ? | ? | 1968.00 | 9.50931 | 0 | 0 |
| BLB | 35139 | *Zonotrichia* | *leucophrys* | 37.6 | -112.8333 | 1970.00 | 6.70134 | 0 | 0 |
| BLB | 35142 | *Zonotrichia* | *leucophrys* | 39.6456 | -111.2603 | 1970.00 | 7.38072 | 0 | 5.37675 |
| BLB | 35143 | *Zonotrichia* | *leucophrys* | 48.6833 | -113.7167 | 1970.00 | 11.58095 | 4.1942 | 0 |
| BLB | 36217 | *Zonotrichia* | *leucophrys* | 37.8994 | -119.2211 | 1975.00 | 10.79638 | 0 | 0 |
| BLB | 37497 | *Zonotrichia* | *leucophrys* | 42.0656 | -124.3083 | 1977.00 | 11.61981 | 0 | 3.93491 |

| | | | | | | | | | |
|---|---|---|---|---|---|---|---|---|---|
| BLB | 37511 | *Zonotrichia* | *leucophrys* | 42.3888 | -122.3306 | 1977.00 | 1.97851 | 0 | 0 |
| BLB | 37792 | *Zonotrichia* | *leucophrys* | 34.4095 | -119.7471 | 1984.00 | 0 | 0 | 0 |
| BLB | 38950 | *Zonotrichia* | *leucophrys* | 37.7698 | -122.4658 | 1984.00 | 14.16514 | 0 | 0 |
| BLB | 39343 | *Zonotrichia* | *leucophrys* | 37.9465 | -119.2063 | 1992.00 | 16.19591 | 0 | 0 |
| BLB | 41320 | *Zonotrichia* | *leucophrys* | 45.2003 | -117.2786 | 1977.00 | 20.5422 | 0 | 0 |
| BLB | 41369 | *Zonotrichia* | *leucophrys* | 34.1006 | -116.8278 | 1977.00 | 27.78624 | 8.2946 | 4.34696 |
| BLB | 42503 | *Zonotrichia* | *leucophrys* | 37.8667 | -122.3 | 1992.00 | 3.97846 | 4.3483 | 0 |
| BLB | 42505 | *Zonotrichia* | *leucophrys* | 37.8667 | -122.3 | 1992.00 | 34.72677 | 0 | 4.37242 |
| BLB | 42835 | *Zonotrichia* | *leucophrys* | ? | ? | ? | 0 | 0 | 4.11112 |

* Individuals marked with asterisks were mis-identified as *Cardinalis sinuatus* in the databases and are now correctly identified as *Cardinalis cardinalis*.
